# Mining Microbial Transcriptomes to Engineer Cell-Based Bacterial Biosensors in Gut-Resident Bacteroidaceae

**DOI:** 10.64898/2026.08.10.744002

**Authors:** Joshua Glazier, David Villegas, Sandra McClure, Joyce Ghali, Jay Fuerte-Stone, Mark Mimee

**Affiliations:** Pritzker School of Molecular Engineering, University of Chicago, 5640 S Ellis Ave, Chicago, 60637, IL, USA; Committee on Microbiology, University of Chicago, 924 E. 57th Street, BSLC R013A, Chicago, 60637, IL, USA; Committee on Molecular Metabolism and Nutrition, University of Chicago, 924 E. 57th Street, BSLC R013A, Chicago, 60637, IL, USA; Department of Microbiology, University of Chicago, 920 E 58th Street, Chicago, 60637, IL, USA

## Abstract

The gastrointestinal tract is rich in metabolic, immune, and microbiome-derived signals that can inform the design of live biotherapeutics and diagnosis of intestinal disorders. Engineered cell-based biosensors can tap into this molecular information and report on their environment, yet their development in gut-resident symbionts has been limited by a lack of validated sensor systems. Here, we present a generalizable pipeline that leverages bacterial transcriptional profiling to identify environment-responsive systems for biosensor engineering. Candidate Sensors Systems (CSSs) mined from healthy, disease, and *in vitro* transcriptomes were assembled into a barcoded library in Bacteroidaceae chassis and screened in high-throughput *in vivo* to identify responsive promoters. A unique Bacteroidales ECF-type sigma factor operon with ties to sphingolipid metabolism and flux was highly responsive in chemically-induced colitis models. The biosensor responded robustly to disease and returned to baseline upon recovery, establishing an *in vivo*-driven strategy for discovering functional biosensors in non-model gut-resident bacteria.

## Introduction

The gastrointestinal tract is a densely interconnected ecosystem where microbes, metabolites, and host immune signals continuously interact to shape physiology, metabolism, and disease susceptibility^1^. Multi-omics studies have detailed how community composition and transcriptional states shift across diet, development, and disease^2–6^, yet the wealth of molecular information encoded in the gut largely remains difficult to measure longitudinally and non-invasiveness. Harnessing information from this biochemically complex environment could inform the diagnosis and monitoring of gastrointestinal diseases for which imaging and blood- or urine-based biomarkers remain insufficient.

Bacteria have evolved specific sensor systems, including transcription factors, two-component signal transduction systems, and alternative sigma factors, to rapidly switch gene expression programs in response to external stimuli ^7–9^. In the mammalian gut, microbial ‘omics studies documented rapid shifts in transcriptional states in response to diet^2^, immune activity^4^, and intestinal inflammation^3,5^, suggesting that host physiology is directly reflected in microbial gene expression. The transcriptional responses of host-associated microbes therefore act as a window into gut biochemistry, encoding real-time information about the intestinal environment.

Since the first proof-of-concept for mercury detection^10^ and foundational gene circuits such as the toggle switch^11^ and repressilator^12^, cell-based biosensors have been designed to co-opt bacterial transcriptional responses to translate biochemical changes into measurable reporter outputs. Unlike conventional diagnostics, bacterial biosensors can colonize the gut and continuously sense disease-relevant biomarkers *in situ*, with proof-of-concept systems already demonstrated for diet^13^, metabolic disorders^14^, pathogenic infections^15–17^, DNA detection^18,19^, inflammatory markers^20–24^ and gastrointestinal bleeding^25^. Beyond diagnostics, sense-and-respond architectures could couple disease detection directly to therapeutic outputs to engineer sentinel cells^26–29^.

Realizing the potential of bacterial biosensors requires both expanding the range of disease-responsive sensors and the repertoire of gut-resident symbiont chassis organisms^29^. Most bacterial biosensors have been constructed in *Escherichia coli* through *ad hoc* adaptation of well-characterized regulatory systems to target a single, pre-defined analyte^27,30^, using bioinformatic^21,31,32^ and high-throughput screening methods^33–35^. Although these approaches are powerful, they are less readily applied to complex disease states for which the relevant molecular signals may be unknown or heterogeneous. Native gut symbionts provide an underexplored source of sensing systems because they are adapted to detect and respond to changes within the intestinal environment. Despite the biocompatibility of gut symbionts and available tools to engineer genera such as *Bacteroides* for gut colonization^36,37^, only a few non-*E. coli* biosensors exist^38–40^. Generalizable strategies are therefore needed to discover disease-responsive biosensors directly from the transcriptional programs of gut-resident bacteria.

Here, we develop a platform to discover and engineer cell-based biosensors in the gut-resident symbionts *Bacteroides thetaiotaomicron* (Bt) and *Phocaeicola vulgatus* (Pv). We use a defined microbial community (DMC) in gnotobiotic mice and RNA-seq to profile member transcriptomes across *in vitro* culture, the healthy gut, and a gnotobiotic dextran sodium sulfate (DSS) colitis mouse model, revealing how gut-resident bacteria adapt and respond to their environment. We identify Candidate Sensor Systems (CSS) via transcriptomic mining and adapt a barcoded reporter high-throughput assay^31^ (BCSeq) to screen candidate’s responses to colitis *in vivo.* Screening hundreds of promoter-based CSSs yielded top candidates that respond to canonical inflammation markers and, unexpectedly, to inflammation-associated sphingolipid synthesis and flux. Finally, we validate a homologous ECF-type sigma factor system in Bt and Pv as engraftable, non-model biosensors responsive to DSS-induced colitis that exhibit low baseline expression, high induced fold change, and dynamic tracking over long-term colonization. This pipeline establishes a generalizable, top-down discovery approach that harnesses gut-resident symbionts for real-time disease monitoring.

## Results

### Mining microbial transcriptional responses to disease states in gnotobiotic mice

For systems-level ecological studies, genome-resolved metagenomics paired with deep metatranscriptomic sequencing is sufficient to learn the families of genes and microbes that respond to an environmental perturbation^41–45^. However, to mine the metagenome for tangible genetic parts, higher genomic resolution is required to pair transcriptional responses with genetic elements retrievable for genetic circuit construction. To reduce the complexity and heterogeneity of the mouse microbiome, we utilized a previously established defined microbial community (DMC)^46^ in gnotobiotic mice as a controlled, physiologically relevant environment for transcriptomic analysis in healthy and disease states (**Figure 1A**). The DMC is comprised of 13 phylogenetically representative human-derived bacterial species that together recapitulate core metabolic capabilities of the human gut microbiome.

**Figure 1.**
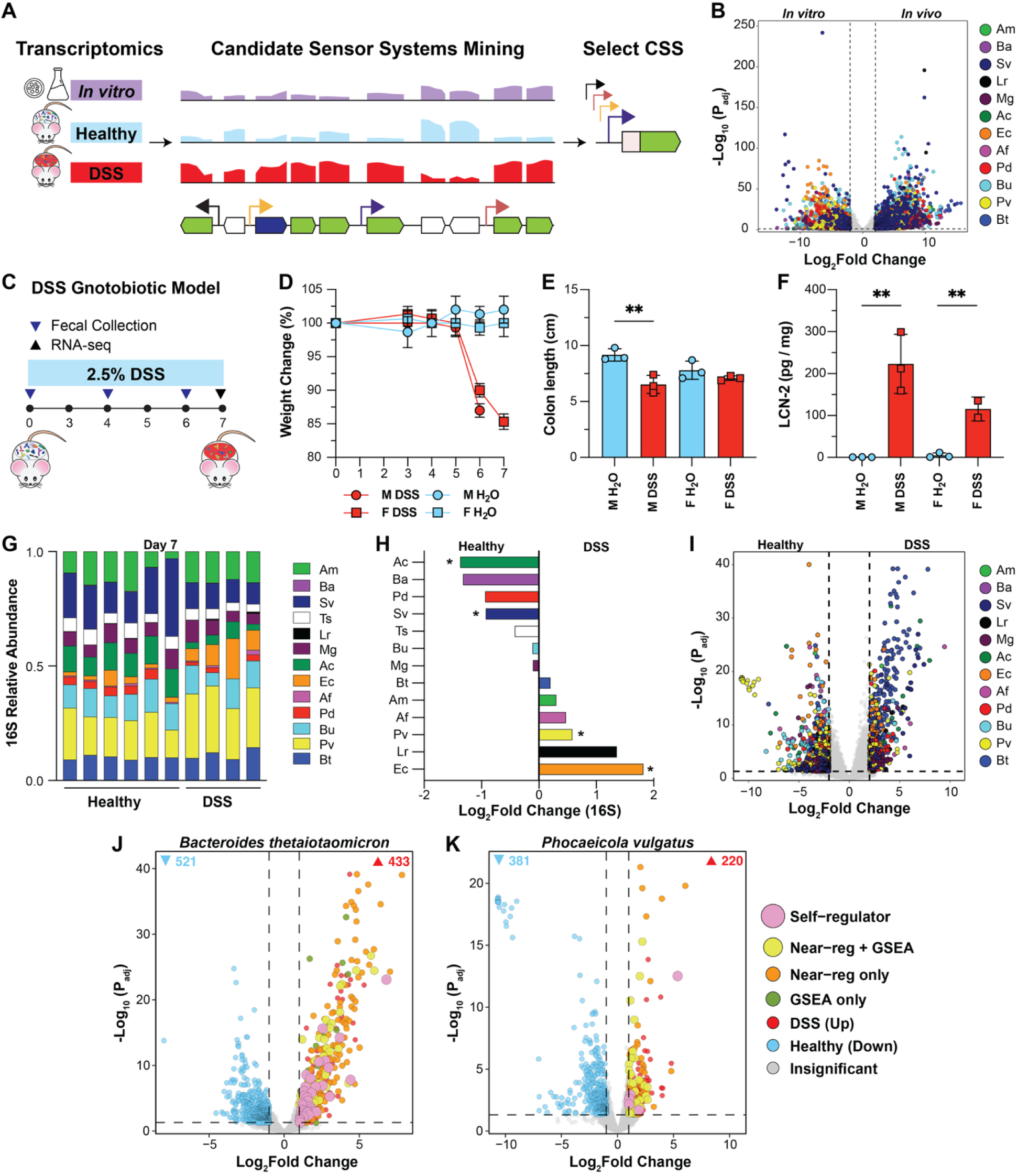
Mining microbial transcriptional responses to disease states in gnotobiotic mice reveals candidate sensor systems of inflammation. (**A**) Schematic of Candidate Sensor System (CSS) mining and selection. RNA-seq analysis of *in vitro,* healthy, and DSS (diseased) conditions is mined for CSS by comparing differential expression, genomic location, operon architecture to define CSS promoters. CSS are selected and engineered to drive reporter expression for quantification of CSS response. (**B**) Species-specific differential expression (DESeq2) across a defined microbial community (DMC) *in vitro* versus *in vivo* cecal contents (n = 3 per condition). Genes with P_adj_ < 0.05 and |Log_2_Fold Change| > 2 are colored by species. (**C**) Schematic of DSS gnotobiotic model wherein 2.5% (w/v) dextran sodium sulfate (DSS) drinking water was administered to DMC-colonized mice on Day 0, with regular drinking water (H_2_O) as a control (n = 6 per condition, equally split male and female). Mice were monitored for 7 days, with fecal collections on Day 0 and Day 4, and euthanized on either Day 6 or 7 for male and female mice, respectively. (**D**) DSS-treated mice lose significantly more weight over time than H_2_O control groups, as a measure of percent change in body weight relative to baseline (Day 0) (p = 2.35e-07, two-way ANOVA with Tukey’s post hoc test). (**E**) Colon length (cm) in male and female mice treated with H_2_O or DSS. DSS treatment significantly reduced colon length in males (p = 0.0096, two tailed Student’s t-test, n = 3 per group) but not in females (p = 0.22, n = 3 per group). (**F**) Lipocalin-2 (LCN-2) concentration (pg/mg cecal contents) in male and female mice treated with H_2_O or DSS. DSS treatment significantly increased LCN-2 in both male and female groups (p = 0.0058 (F), p = 0.0054 (M), two tailed Student’s t-test, n = 3 per group). (**G**) Relative abundance of DMC from 16S rRNA sequencing across Healthy (H_2_O, n = 6) and DSS (n = 4) groups from Day 7 cecal contents (Day 6 for M DSS). (**H**) Differential abundance (Log_2_fold change, 16S) of DMC community members from Day 7 cecal, DSS versus Healthy. *Phocaeicola vulgatus*, and *Escherichia coli* significantly increased (p < 0.05, Kruskal-Wallis, Mann-Whitney U), and *Anaerostipes caccae, Subdoligranulum variabile* significantly decreased (p < 0.05, Kruskal-Wallis, Mann-Whitney U). (**I**) Species-specific differential expression (DESeq2) across the Defined Microbial Community (DMC) Healthy versus DSS cecal contents (n = 6 Healthy, n = 5 DSS). Genes with P_adj_ < 0.05 and |Log_2_Fold Change| > 1 are colored by species. (**J-K**) *Bacteroides thetaiotaomicron* and *Phocaeicola vulgatus* individual differential expression, genes colored according to SSMiner CSS classification. Abbreviations: Bf, *Bacteroides fragilis*; Cc, *Coprococcus comes*; Am, *Akkermansia muciniphila*; Ba, *Bifidobacterium adolescentis*; Sv, *Subdoligranulum variabile*; Ts, *Turicibacter sanguinis*; Lr, *Limosilactobacillus reuteri*; Mg, *Mediterraneibacter gnavus*; Ac, *Anaerostipes caccae*; Ec, *Escherichia coli*; Af, *Alistipes finegoldii*; Pd, *Parabacteroides distasonis*; Bu, *Bacteroides uniformis*; Pv, *Phocaeicola vulgatus*; Bt, *Bacteroides thetaiotaomicron*.

To establish the feasibility of metatranscriptomic profiling and CSS mining in the DMC in gnotobiotic mice, transcriptional profiles were compared between *in vitro* mid-log growth and multiple locations of a healthy intestinal tract (**Figure S1**). Fecal sampling at the cecum and colon showed similar compositions and transcriptomes (**Figure S1A-C)**, while ileum reads were dominated by *E. coli*, likely due to lower biomass and a more oxygenated environment. Cecum samples were analyzed going forward for consistency when comparing *in vitro,* healthy and disease transcriptomes. Overall, comparison of *in vitro* cultures and the *in vivo* DMC community showed extensive differential expression across all community members (**Figure 1B)**. Broadly, carbohydrate utilization genes were upregulated *in vivo*, reflecting the more diverse nutrient landscape of the gut, while ribosomal and electron transport chain genes were downregulated, consistent with a reduced *in vivo* growth rate^47^ and a shift towards fermentative metabolic strategies (**Table S8**).

Next, as a disease model for biosensor discovery, we implemented a chemical model of colitis by supplementing drinking water with dextran sodium sulfate (DSS). Inflammation causes well-documented, multifaceted changes to the gut biochemical landscape, including shifts in metabolites and SCFAs^48^, alterations in host immune responses, and barrier integrity failure^3,5,6,49^. These changes provide numerous potential signals for microbial sensing and represent a significant pool of biochemical inducers that trigger rapid transcriptional shifts in resident bacteria^3–5^. Directly mining these transcriptional responses harnesses natural bacterial sensor systems for biosensor design and bypasses the need for explorative metabolomics to identify new biomarkers. Gnotobiotic mice colonized with the DMC were treated with 2.5% DSS in drinking water for 7 days, and cecal contents harvested for RNA-seq analysis on Day 7 (**Figure 1C**). As expected, DSS-treated mice lost significantly more weight than water controls (**Figure 1D**, p = 2.3514e-07, Two-way ANOVA, with Tukey’s post hoc test) and displayed signs of illness including lethargy, reduced grooming, and bloody stool, indicating substantial gut inflammation. DSS-treated mice also had shortened colons (**Figure 1E**) and increased fecal lipocalin-2 concentrations on Day 6/7 (**Figure 1F**). DSS treatment induced compositional shifts in the DMC by Day 7, with increases in *E. coli* and *P. vulgatus* and depletion of butyrate-producing obligate anaerobes such as *Anaerostipes caccae* and *Subdoligranulum variabile* (**Figure 1G-H, Figure S2A-C**). These taxonomic shifts align with established patterns in colitis: Pseudomonadota increase, Bacteroidota and Bacillota trade off, except for some putative pathobionts, such as *Mediterraneibacter gnavus*^5,6^.

Beyond composition shifts, transcriptional profiling revealed how individual community members respond to colitis. RNA-seq showed clear separation of DSS and water groups by PCA of variance-stabilizing transformed counts (**Figure S2E**) and thousands of significant differentially expressed genes (DESeq2, log₂FC > 1, P_adj_ < 0.05) across the community (**Figure 1I, Figure S2F, S3)**. We observed few differences in low-abundance members (*Limosilactobacillus reuteri, Bifidobacterium adolescentis*) differential expression due to read depth. Differential expression dynamics varied species-to-species in the community. Members with known aerotolerance and increased or stable abundance in DSS samples, such as *Bacteroides thetaiotaomicron, Phocaeicola vulgatus,* and *Escherichia coli*, accounted for most of the upregulated differential expression. Gene Set Enrichment Analysis (GSEA) of SEED subclass, SEED subsystems, and available KEGG pathways revealed a community under broad physiological stress and metabolic remodeling (**Figure S4).** The most widespread response was downregulation of carbohydrate utilization, including Bacteroidaceae PULs and sugar utilization pathways across phyla, a dynamic that reflects a broad metabolic shift toward survival in the inflamed gut and has been previously reported in metagenomic surveys of IBD patients^50,51^. The community-wide impact of DSS-induced stress was also apparent; upregulated subsystems, including an aerotolerance operon (*Bacteroides uniformis),* protein chaperones and folding (Bt, Pv, *Parabacteroides distasonis, Mediterraneibacter gnavus)*, stress response (Bt, *Parabacteroides distasonis*), the PHO regulon (Bt, Pv), and fatty acid biosynthesis and membrane repair pathways (Pv, *Bacteroides uniformis*) are consistent with broad physiological stress across community members. Notably, *Akkermansia muciniphila* remained largely stable, with few DEGs or enriched pathways, suggesting resilience in both growth and transcriptional response.

### Bacteroidaceae spp. are strong candidate chassis organisms for biosensor development

Due to their substantial transcriptional responses across all conditions, community prevalence and stability, capacity for controllable engraftment^52^, and robust genetic tools^53,54^, Bt and Pv were selected as chassis organisms for transcriptomic mining to identify CSSs. *In vivo* versus *in vitro* transcriptomics primarily provided confidence in the transcriptional approach for CSS mining. The analysis showed Bt and Pv as robust responders to the distinct growth environments and recapitulated known Bacteroidaceae functions in the gut, such as activation of PUL operons (**Figure S1D-F)** involved in degrading dietary fibers and plant-derived substrates^55–57^, a well-documented response to nutrient availability^58–61^. Applied to the DSS colitis model, both species acutely responded to the disease conditions and mounted metabolic, stress-response, and membrane remodeling programs. With a high number of differentially expressed genes, and many tied to predicted inflammation response systems, Bt and Pv are well-suited for CSS discovery and screening.

With Bt and Pv identified as suitable chassis, we developed a custom computational pipeline (SSMiner), inspired by TFBMiner^62^, to systematically mine CSS candidates from their transcriptomes. SSMiner integrated differential expression, GSEA, regulatory context, and operon structure to rank and prioritize promoter candidates (**Materials and Methods)**. Candidates were binned by regulatory class (self-regulator or near-regulator) and by GSEA enrichment categories (**Figure 1J-K**). From 433 (Bt) and 220 (Pv) upregulated genes, SSMiner ranked 333 Bt and 162 Pv candidate promoters, using operon mapping to define CSS structure. Top-ranked candidates were those in multi-gene operons with high differential expression across all operon members and either self-regulation or a defined proximal regulator. Incorporating GSEA pathway information further improved CSS selection and Pool design in later iterations. CSSs were then selected iteratively for promoter synthesis and screening through a Design-Test-Build-Learn cycle (**Figure 2A**).

**Figure 2.**
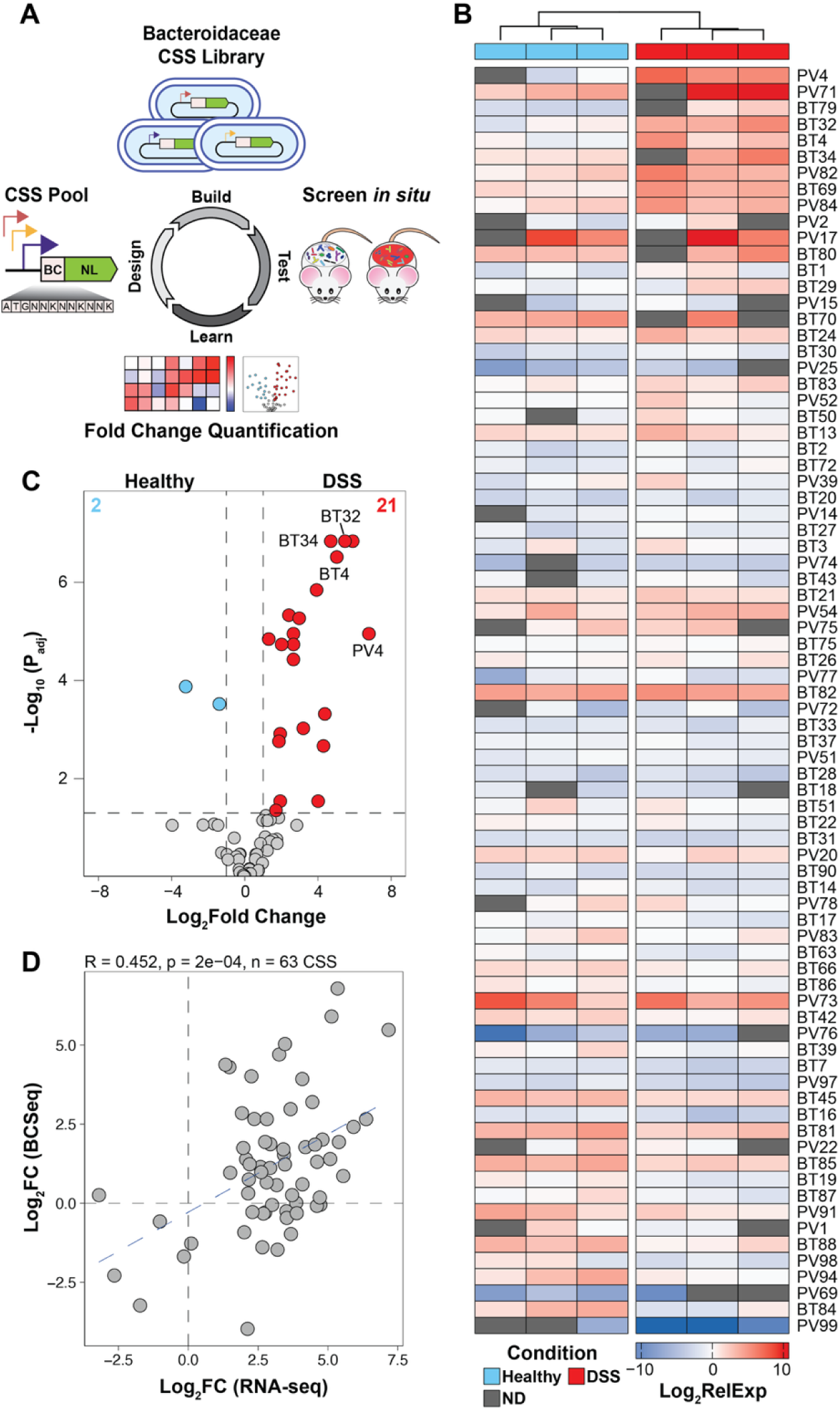
Design-Build-Test-Learn iterative screening of pooled Bacteroidaceae candidate sensor systems in vivo identifies putative gut inflammation responsive-regulatory elements. (**A**) Schematic of Design-Build-Test-Learn approach to screening CSSs. CSS Promoters are engineered behind a 9-bp barcoded (BC) Nanoluciferase (NL) reporter gene (Design). CSS circuits are integrated into *Bacteroidaceae* (Bt and Pv) and pooled to create CSS Pools for screening (Build). CSS Pools are engrafted into mice and screened *in vivo* during DSS and Healthy conditions (Test). Individual CSS responses to DSS-induced conditions are quantified using Barcode Sequencing (BCSeq) to determine successful candidates and design the next pool (Learn). (**B**) Heatmap of BCSeq Relative Expression (Log_2_RelExp) for CSS Pool3 in DSS and Healthy conditions (n = 3, per condition). Rows are sorted top to bottom by RelExp fold change (DSS to Healthy) and hierarchal clustering determined column order. Barcodes not detected (ND) are colored gray. (**C**) MPRAnalyze results of CSS Pool3 fold change in DSS versus Healthy. 21 CSS were significantly upregulated (Log_2_Fold Change > 1, P_adj_ < 0.05, Benjamini-Hochberg) and 2 CSS significantly downregulated (Log_2_Fold Change < −1, P_adj_ < 0.05, Benjamini-Hochberg). (**D**) Comparison of BCSeq (Log_2_FC) and RNA-seq results (Log2FC) for CSS Pool3 by Pearson correlation (R = 0.452, p = 2e-04).

### A novel method for high throughput in vivo pooled screening of candidate sensor systems

Screening each CSS individually *in vivo* would require hundreds of mice, making individual testing impractical. Conversely, while *in vitro* screening could narrow the candidate pool, it would rely on selecting putative disease-associated stimuli and biomarkers *a priori* and testing them in simplified conditions that do not recapitulate the complex features of the gut environment, including microbe- and host-microbe interactions, spatial variation along the intestinal tract, and simultaneous exposure to multiple signals, particularly during disease. Furthermore, since baseline microbial transcriptional profiles vary drastically between *in vivo* and *in vitro* states (**Figure 1B**), sensors that appear inactive, constitutive, or strongly inducible *in vitro* may behave differently *in vivo*. We therefore sought to develop a pooled, barcoded approach that could simultaneously evaluate hundreds of CSS directly within the relevant intestinal environment.

To bridge this gap, we developed a high-throughput, barcode-based platform for functional biosensor screening, adapted from a previously described pooled method for gene expression profiling^31^. Endogenous promoter regions of selected CSSs were registered with a CSS ID according to chassis and operon architecture (i.e. BT1 or PV1) and cloned upstream of a barcoded NanoLuc (NL) reporter within a pNBU1 *Bacteroides* integration vector ^63^. Each promoter was assigned multiple unique 9 base pair barcodes inserted immediately after the NL start codon, enabling independent measurements of promoter activity by quantifying barcodes in transcripts. The barcoded promoter pool was transformed into *E. coli* S17 λ*pir* and transferred by conjugation into the Bacteroidaeceae species from which each promoter was originally derived. Using promoters endogenous to the host organism ensures the presence of necessary regulators and establishes a characterized baseline for promoter activity. The resultant Bt and Pv CSS pools were then assayed in different conditions, after which RNA and genomic DNA was extracted in parallel from each bacterial population. Barcode-containing transcripts were selectively reverse-transcribed using an NL-specific primer and prepared for BCSeq through adapter ligation and amplification. RNA barcode abundance was then normalized to the abundance of the corresponding barcode in genomic DNA, thereby controlling for differences in strain representation within the pooled population. This BCSeq workflow generated a normalized expression measurement for each promoter and enabled the simultaneous evaluation of hundreds of CSSs in a single experiment.

### High-throughput discovery of colitis-responsive candidate sensor systems

Using an iterative design-test-build-learn methodology, we engineered CSSs into consecutive pools for high-throughput *in vivo* screening. In our initial pilot library (Pool1), 52 total selected CSSs for Bt and Pv were individually amplified from genomic DNA, cloned into the Bacteroidaceae integrative vector with random barcodes, and pooled prior to engraftment into germ-free mice alongside the remaining members of the DMC. Each CSS was confirmed for reporter expression prior to incorporation into the pool. Additionally, since the inserted barcode altered the primary amino acid sequence of NL, we measured transcript abundance and luminescence for a representative CSS in each chassis. While output luminescence varied up to 10-fold across different barcodes, transcription of the NL reporter did not differ, suggesting the method can be used to measure promoter transcriptional activity (**Figure S5**).

Following Pool1 engraftment, DSS administration induced weight loss, colon shortening, and elevated lipocalin-2 levels, consistent with the RNA-seq experiments (**Figure S6A-D)**. Notably, there was no significant increase in luminescence from fecal contents from the pooled CSSs populations (**Figure S6E),** underscoring the need for barcode-level resolution to measure promoter activity. Importantly, CSSs with more than one barcode were highly correlated across both conditions (**Figure S6F)**, suggesting reproducibility with the method. To determine CSS response to colitis, we applied two complementary analytical approaches. First, we calculated median-normalized relative expression (RelExp) per CSS per sample to provide sample-level visualization of promoter activity and absolute promoter strength across conditions (**Figure S6G**). Next, we applied MPRAnalyze^64^, a hierarchical negative binomial model, to calculate fold changes from raw counts, accounting for experimental variability between library preparations, barcodes, engraftment differences, and unbalanced group sizes from gnotobiotic mouse availability. In Pool1, 10 CSS significant increased response to colitis (**Figure S6H)** and comparisons between the measured fold changes of BCSeq and RNA-seq showed positive correlation (R =0.415; p=0.0097, Pearson correlation) **(Figure S6I)**. Candidates BT32 and PV4 emerged as the two most robust responders. Altogether, the functional biosensor screening platform allowed for pooled profiling of CSSs for the identification of disease-responsive promoters.

While Pool1 established the viability of the CSS BCSeq pipeline, individual cloning and retroactive barcode assignment proved to be labor intensive and inefficient. To scale to a more high-throughput approach, we designed and engineered an expanded pool comprising 67 CSSs per chassis from our SSMiner analysis using synthesized oligopools. Each CSS was designed with two pre-assigned barcodes (134 promoters, 268 barcodes total), serving as internal technical replicates for quality control and guarding against colonization dropouts observed in Pool1. Following integration into their respective hosts, the libraries were pooled for engraftment. DNA BCSeq of the initial plasmid libraries detected 258 of 268 barcodes and 133 of 134 CSS in a near-normal distribution for each species, although conjugation into Pv showed a slight skewing attributed to lower mating efficiencies during library preparation (**Figure S7F-G**). Pool2 was screened in healthy and DSS mice (**Figure S7A-E)**, alongside an additional *in vitro* culture screening to define baseline CSS activity, confirm transcriptional function, and ensure each CSS was sampled without engraftment bottlenecks. These control experiments validated the quantitative accuracy of BCSeq, demonstrating tight correlations within individual samples and chassis species (**Figure S8, Figure S9A-B)**. Quantitative analysis revealed 18 significantly upregulated CSS from DSS-induced colitis (**Figure S9D**; Log_2_FC > 1, P_adj_ < 0.05). Comparison of BCSeq and RNA-seq results showed significant correlation between CSS Log₂FC and median RNA-seq Log₂FC values for the CSS operon (**Figure S9E**; R =0.564; p= 3.1e-07, Pearson correlation). Importantly, of the 18 upregulated CSS, BT32 and PV4 again were the top induced sensors, reproducing the observations in Pool1 screening.

Despite successful engraftment alongside the DMC and detection *in vitro*, a subset of Bt and Pv candidates in Pool2 were transcriptionally inactive. Re-analysis of Pool2 CSSs revealed missing transcriptional start sites^60^ for many inactive CSS, likely due to the assigned promoter size during Pool2 oligopool synthesis. Additionally, certain CSS candidates with strong baseline or disease-induced expression dominated the BCSeq read counts, reducing the resolution for weaker promoters. We used the structural insights and performance metrics from Pool1 and Pool2 to re-design a third iteration comprised of 100 CSS candidates (50 in Bt and 50 in Pv), each with 2 barcodes (Pool3). To optimize this library, we emphasized longer promoter lengths during synthesis, focused on highly active GSEA pathways, and streamlined overall pool complexity. Specifically, we carried forward the most promising CSS from previous pools, eliminated measured non-sensing candidates and those with excessive baseline activity, and integrated a small subset of downregulated CSSs and housekeeping genes to anchor the dynamic range of the assay. DNA BCSeq detected 184 of 200 barcodes and 96 out of 100 Pool3 CSS in the final engineered pool. Overall, both chassis libraries showed a near-normal distribution, with a noticeable reduction in Pv CSS skewing (**Figure S10E**). Pool3 was engrafted with the DMC into gnotobiotic mice, treated with DSS to induce colitis, and measured via BCSeq (**Figure S10, Figure 2B)**. Pool3 showed a greater number of CSS induced in colitis conditions (**Figure 2C)**, with 16 and 5 activated in Bt and Pv, compared to 15 and 1 in Pool2, respectively. The global correlation of Pool3 BCSeq to bulk RNA-seq differential expression was significant (**Figure 2D**; R =0.452; p= 2e-04, Pearson correlation). Additionally, Pool3 yielded more predicted CSS having a positive fold change (**Figure 2C**; 21, 33.3% of measured CSS), as compared to Pool1 (10, 25.6%) and Pool2 (18, 24.6%). As with Pools 1 and 2, BT32 and PV4 were validated among the most powerful and reproducible responders, alongside several new candidates (**Figure 2B-C**). Six previously untested CSS were significantly upregulated (BT79, BT80, BT83, PV77, PV82, PV84), highlighting improved mining of transcriptomic data in Pool3, while multiple other CSS (BT20, BT27, BT29, BT4) were now validated across multiple Pools. Previously tested CSS BT30 and BT34 were among most highly induced CSS in Pool3 despite showing no induction in earlier pools, suggesting that expanded promoter size improved the screen. Pool 3 design improvements also yielded more upregulated CSS in Pv (5), a validated downregulated candidate (BT84), and housekeeping promoters that were detected but not differentially expressed (BT87, PV94). Incorporating a Design-Build-Test-Learn framework markedly improved the BCSeq screening process, establishing a powerful, scalable *in vivo* platform for candidate biosensor discovery.

### Most top colitis-responsive candidate sensor systems are induced by canonical inflammation stressors

From the almost 200 total promoters screened via BCSeq across Bt and Pv, we selected 10 top candidates for further individual validation as biosensors of DSS-induced colitis (**Table S16**). Each CSS was individually cloned to remove barcodes that may impact NL translation and reintroduced into its host chassis. We then tested each CSS in *in vitro* culture assays to evaluate responses to a panel of canonical inflammation stressors, including oxidative stress, low pH, and altered bile acid exposure. We also assayed DSS directly (1% w/v) as a negative control to confirm that *in vivo* CSS induction reflects a disease-relevant response rather than a direct interaction with the DSS molecule.

The CSSs exhibited distinct, but partially overlapping response profiles *in vitro.* Exposing anaerobic, mid-log cultures to a two-hour oxygen shock via aerobic shaking stimulated reporter expression across half of the candidates, with several CSSs showing up to 10-fold induction (**Figure 3A**). Among the top oxygen-responsive systems were BT14 and BT34, which drive expression of thioredoxins (*BT_0549* and *BT_0218*/*BT_0219* respectively), and have functional ties to oxidative stress. Because several CSSs responded to oxygen exposure, we investigated whether their activity depended on OxyR, a major regulator of oxidative-stress responses in Bt. We deleted *oxyR* (*BT_4716*) in Bt and integrated five oxidative stress-responsive CSSs into the mutant strain. Under anaerobic conditions, basal BT34 expression increased over 1,000-fold in the *oxyR* mutant (**Figure S11B**), providing genetic evidence that OxyR represses this promoter in the absence of oxidative stress. A second subset of candidates, including BT14, BT28, BT29, and BT34, was responsive to growth in acidified media (BHIS, pH = 5.3). Bile acid exposure activated a third subset of candidates, including BT3 (*BT_0884-BT_0886*; efflux system for antibiotic and toxins), BT4 (*BT_0954-BT_0957*; antimicrobial resistance), and BT27 (*BT_2567-BT_2568*; acid resistance). BT34 was notably induced in all three conditions, suggesting it responds to a shared stress-associated signal. In contrast, direct *in vitro* addition of 1% DSS to cultures failed to induce substantial reporter gene expression across the top candidates, indicating that their disease-associated induction was not attributable to direct sensing of DSS.

**Figure 3.**
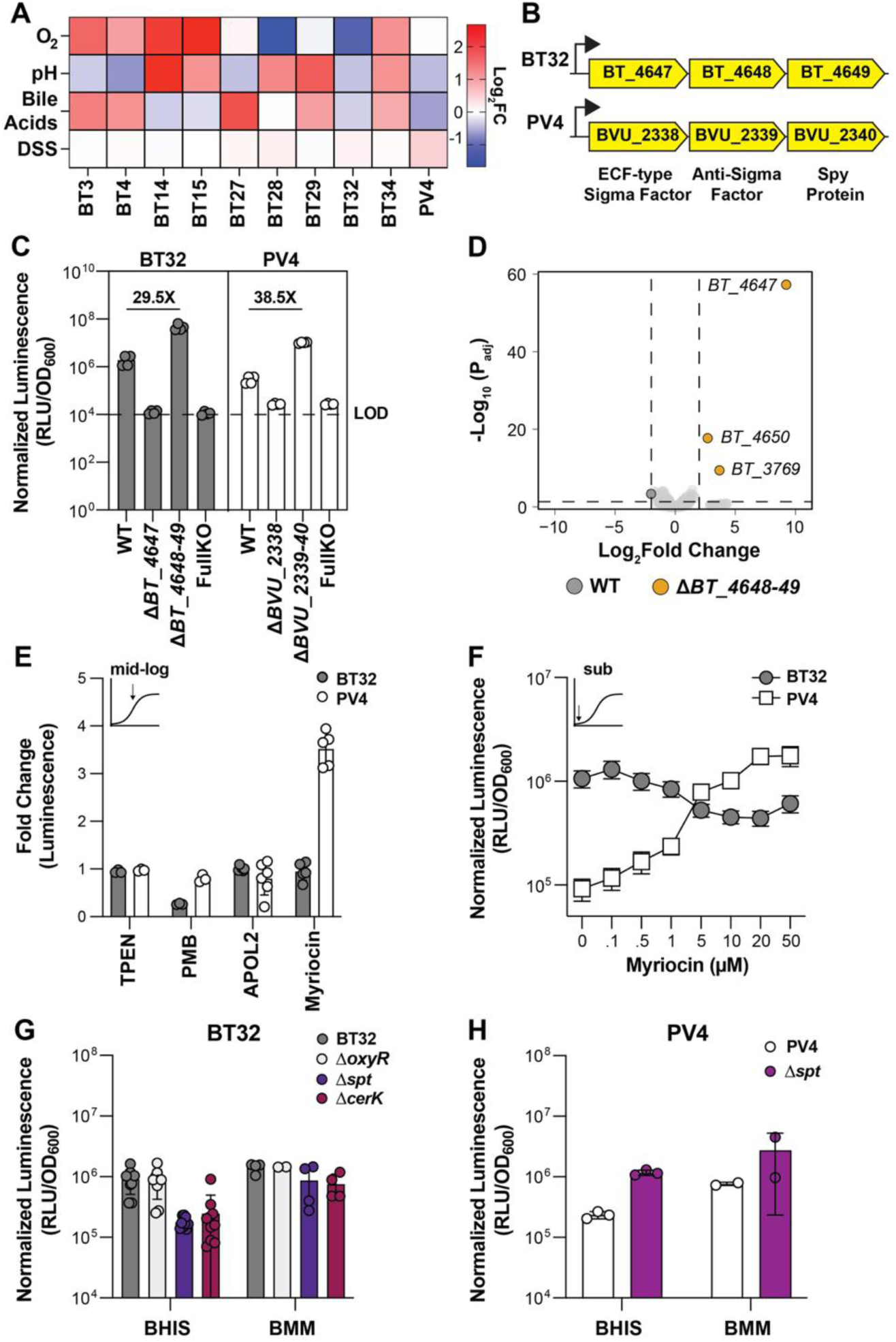
Top colitis-responsive candidates respond to canonical inflammation stressors in vitro, while homologous CSSs BT32 and PV4 drive self-regulating ECF-type sigma factor operons responsive to changes in sphingolipid flux. (**A**) Fold change response (Log_2_FC) of top CSSs (Table S16) to canonical inflammation stressors, oxidative stress (O_2_, via aerobic shaking), acidic conditions (pH = 5.3, BHIS media), and bile acids (Ox bile acids, 0.03% w/v in BHIS media). Dextran sodium sulfate (DSS, 1% w/v) was tested to confirm limited response from each CSS. CSSs were induced at mid-log growth in BHIS media, then sampled 2 hours post induction to calculate fold changes by comparing normalized luminescence values (RLU/OD_600_) of induced vs uninduced control cultures. (**B**) Three-gene operon architecture of CSS BT32 and PV4. Gene annotations determined by SeqHub. (**C**) Expression of BT32 and PV4 NL reporter after integration of CSS circuit into knockout mutant strains of Bt and Pv (Figure S12A, Table S5). Deletion of the anti-sigma factor and Spy protein resulted in 29-38 fold increase in CSS expression in Bt and Pv. Deletion of ECF-type sigma factor or full knockout (FullKO) in Bt and Pv reduced reporter expression to the limit of detection (LOD). Expression measured as normalized luminescence (RLU/OD_600_). (**D**) Differential expression of *ΔBT_4648-49* versus WT Bt. 3 genes (*BT_4647, BT_4650, BT_3769*) were significantly upregulated (Log_2_Fold Change > 2, P_adj_ < 0.05). (**E**) Response (Fold change in Normalized Luminescence) of BT32 and PV4 to mid-log growth induction with membrane stressors *in vitro*. TPEN (10 μM), Polymyxin B (PMB, 50 μg/mL), and Apolipoprotein2 (APOL2, 10 μg/mL in BHI media) failed to induce BT32 and PV4 when compared to control conditions. Myriocin (20 μM) induced a response in PV4. n = 3-5 per condition. (**F**) Dose-dependent response of BT32 and PV4 to increasing concentrations of myriocin *in vitro* when induced from overnight subculture. (**G**) BT32 promoter expression in WT Bt and *oxyR* (*BT_4716), spt* (*BT_0870) and cerK* (*BT_0871)* deletion mutant Bt strains for overnight cultures in multiple media types (BHIS, B-MM). In both media types, deletion of *oxyR* has no effect on expression, while deletion of *spt* and *cerK* reduces BT32 expression. (**H**) PV4 promoter expression in WT Pv and *spt* (*BVU_3166*) deletion mutant Pv strain for overnight cultures in multiple media types (BHIS, B-MM). In both media types, deletion of *spt* increases PV4 expression.

Successful stimulation of these candidates by canonical inflammation stressors confirmed that the BCSeq screen effectively selected for CSS with functional ties to *in vivo* inflammatory environments. In particular, the activation of a general thioredoxin system (BT34) across all tested stressors, paired with its dramatic repression in the *oxyR* mutant, served as a robust internal positive control for the platform. In contrast, the two highest-ranked BCSeq candidates, BT32 and PV4, did not respond to any canonical inflammatory stressors. While PV4 was modestly induced by 1% DSS, additional testing in alternative media (BHI, B-MM) confirmed minimal *in vitro* induction for both BT32 and PV4 (**Figure S11A**). Their strong and reproducible activation during colitis, despite the absence of an identifiable inducer under these simplified conditions, suggests that they respond to a feature of the intestinal environment not captured by the canonical stress assays. We therefore selected BT32 and PV4 for deeper characterization.

### The BT32 and PV4 candidate sensor systems control expression of an ECF-sigma factor regulatory system

Across RNA-seq profiling and BCSeq screens, candidates BT32 (*BT_4647-BT_4649*) and PV4 (*BVU_2338-2340*, recently described as *cirYZ* in the isolated human donor strain PvCL10^65^) consistently emerged as the strongest candidate biosensors, showing high expression during DSS-induced inflammation and low baseline expression in healthy mice. BT32 and PV4 drive expression of a homologous three-gene operon, consisting of an ECF-type sigma factor, a membrane-associated anti-sigma factor, and predicted Spy-like chaperone protein (**Figure 3B)**. ECF-type sigma factors are bacterial signal-transduction systems that regulate promoter activity in response to extracellular and envelope-associated stress ^66–68^. Bacteroidales genomes are uniquely enriched in ECF sigma factors; among these, *BT_4647* and *BVU_2338* belong to the ECF type-3 subfamily, which is nearly exclusive to Bacteroidales with a specific −33/-7 motif and 17-bp gap in associated promoter regions^67^ (**Figure S12D)**. Structural modeling (AlphaFold3^69^ and SeqHub^70^) of the remaining two genes suggested conserved roles, annotating the second gene as an anti-sigma factor with a transmembrane domain, and the third as a periplasmic Spy-like chaperone (**Figure S12F)**. Together, these features propose a model where the membrane associated anti-sigma factor sequesters the ECF under basal conditions, resulting in a low baseline promoter activity until signal detection at the membrane releases or activates the sigma factor to drive feed-forward transcription of the full operon, including the secreted Spy-like chaperone that may interact with misfolded periplasmic proteins^71^. This predicted architecture suggests that BT32 and PV4 may respond to canonical ECF sigma factor inputs, such as membrane stress, metal availability, or cationic antimicrobial peptides^72–75^.

Amino acid sequence alignment of the operons revealed at least 55% shared homology for each gene, with the ECF-type sigma factor the most conserved at 73%. We confirmed that the BT32 and PV4 promoters are functional in both Bt and Pv, though expression level is host-dependent (**Figure S12E)**. Despite these similarities, BT32 and PV4 occupy different genomic loci with distinct neighboring genes. To determine whether this operon structure is conserved more broadly, the BT32 and PV4 operons were analyzed through a BLAST-base synteny analysis using the biological agent Biomni^76^. Of the 35 named species carrying all three genes, 34 maintained strict operon synteny with a conserved gene order. The operon is distributed throughout Bacteroidales, with syntenic hits in *Bacteroides*, *Phocaeicola*, *Mediterranea, Segatella*, and *Prevotella* species (**Figure S13**), suggesting a conserved adaptive mechanism throughout Bacteroidales.

Next, to test the predicted regulation mechanism, we constructed chromosomal deletion mutants of the ECF-type sigma factor (*ΔBT_4647/ ΔBVU_2338*), anti-sigma factor and Spy-like protein (*ΔBT_4648-49/ΔBVU_2339-40*) and full operon in Bt and Pv (**Figure 12A**). Deletion strains were conjugated with their cognate BT32- and PV4 NL-expression vector to use reporter expression as a proxy for promoter activity. In both species, *ΔBT_4647/ ΔBVU_2338* and full KO abrogated baseline promoter activity, with expression at media-only controls levels (**Figure 3C).** In *ΔBT_4648-49/ΔBVU_2339-40* mutants, deletion of the anti-sigma factor and Spy-like protein increased promoter expression approximately 30–40-fold, suggesting that the ECF is inhibited by one of the two downstream proteins. We next performed transcriptomic analysis of wild-type Bt and the deletion strains *ΔBT_4647* (**Figure S12C)** and *ΔBT_4648-49* (**Figure 3D)** to explore the breadth of the ECF sigma factor regulon. Principal component analysis of DESeq2-transformed data showed no separation by genotype (**Figure S12B**), and differential expression analysis between *ΔBT_4648-49* and WT had 3 significant genome-wide transcriptional changes (**Figure 3D**), including *BT_4647* (as expected), *BT_4650* (a voltage-gated chloride channel immediately downstream of BT32 operon), and *BT_3769* (a hypothetical protein structurally similar to a periplasmic Spy protein). The *BT_4647* ECF sigma factor regulates a narrow set of genes in response to colitis-associated stress.

### The BT32 and PV4 candidate sensor systems respond to select changes in sphingolipid flux

Given the predicted structural annotations of an ECF-type sigma factor and a Spy-like chaperone, we hypothesized that BT32 and PV4 respond to envelope stress associated with periplasmic misfolded proteins. Bile acid and acidic pH induction at mid-log growth did not induce expression (**Figure 3A**), nor did subculturing in the presence of heat or acidified media did not activate either promoter (**Figure S15A**). Neither sensor activated in response to metal chelation via TPEN (**Figure 3E**), despite structural-homology analyses suggesting potential connections to heavy-metal sensing. Mid-log induction with polymyxin B treatment reduced promoter expression in Bt in a dose-dependent manner (**Figure 3E**, **Figure S15B**), but did not activate PV4, although previous TnSeq studies implicated the operon in polymyxin B resistance in *P. vulgatus*^77^. Thus, BT32 and PV4 were not activated by several canonical forms of envelope, thermal, or acid stress, but BT32 did respond to cationic peptide treatment.

To screen a broader range of conditions, we repurposed Biolog Phenotype MicroArray plates as high-throughput inducer exploration assays. We selected plates P4 (phosphorus and sulfur utilization) and P9 (osmotic and ionic stress) to test inflammation-relevant compounds and membrane stressors (**Figure S14)**. Ethylene glycol (EG) at concentrations up to 20% was the most effective inducer (**Figure S14G)** while increasing sodium benzoate concentrations repressed BT32 and PV4 expression (**Figure S14F)** potentially acting as a membrane stabilizer. Together, these results indicate that BT32 and PV4 are not broadly activated by canonical inflammatory or envelope stressors but can be modulated by select chemical perturbations with potential effects on envelope physiology.

Because neither the targeted stress assays nor the broader chemical screen identified an obvious physiological inducer, we next examined prior studies for pathways associated with the BT32 locus. Recent reports correlated upregulation of the BT32 operon to outer membrane vesicle (OMV) biogenesis and sphingolipid production^78,79^. In a proposed model, host-derived apolipoprotein 9a/2 (APOL9a/2) binds to the *Bacteroides* outer membrane sphingolipids, driving OMV formation and release as part of a host-microbe crosstalk pathway regulating gut immunity^79^. Separately, BT32 operon expression was linked to the deletion of serine palmitoyltransferase (SPT)^78^, the enzyme that catalyzes the first committed step in bacterial sphingolipid biosynthesis and there is a previously reported role of sphingolipids in polymyxin B resistance^80^. Together, these observations suggested that BT32 and its homologous PV4 system might respond to perturbations in sphingolipid production or membrane lipid homeostasis.

Although upregulation of the BT32 operon was previously reported during *in vitro* APOL2 treatment^79^, direct addition of recombinant APOL2 to BHI cultures failed to increase reporter expression for either BT32 or PV4 in our assay conditions (**Figure 3E**). We next treated mid-log BT32 and PV4 reporter strains with myriocin, a chemical SPT inhibitor^81,82^. Following treatment in BHIS media, PV4 expression increased nearly 4-fold, whereas BT32 was unchanged (**Figure 3E)**. We subsequently tested a broader range of myriocin concentrations, growth phases, and culture media. When the biosensor strains were subcultured in the presence of myriocin, PV4 activity increased, whereas BT32 activity decreased, with both systems exhibiting divergent, but dose-dependent responses (**Figure 3F, Figure S15C-D)**. Thus, pharmacological inhibition of sphingolipid biosynthesis consistently modulated both sensor systems, prompting us to investigate their relationship to sphingolipid metabolism in greater detail.

To genetically validate the connection between sphingolipid metabolism and these ECF-type sigma factor operons, we deleted the *spt* gene in both Bt (*BT_0870*) and *Pv* (*BVU_3166*). To test distinct steps in the sphingolipid synthesis pathway, we additionally deleted the *cerK* gene (*BT_0871*) in Bt, which encodes the enzyme responsible for synthesizing ceramide-1-phosphate, the lipid to which APOL9a/2 binds^79^. Integration of BT32-NL and PV4-NL reporter circuits into these mutant strains showed promoter expression changes that mirrored the myriocin induced phenotype (**Figure 3G-H)**. Bt *ΔoxyR* was included as well as a negative control, confirming that changes in expression were linked the sphingolipid disruption, and not general stress sensing. Deletion of the *spt* and *cerK* genes in Bt, and *spt* in Pv, affected growth rates and emulated myriocin-treated strains normalized reporter luminescence (**Figure S15E-F, H-I**). SPT deletion removed all response to myriocin in Bt, and blunted the fold change increase in Pv, relative to the wild-type controls (**Figure S15G, J).** Together, these *in vitro* investigations into the mechanism of BT32 and PV4 response *in vivo* point to an associated with endogenous sphingolipid synthesis and flux.

### In vivo validation of BT32 and PV4 biosensors across colitis models

Based on their robust performance in *in vivo* screening and unique potential biological function, biosensors BT32 and PV4 were advanced to individual *in vivo* functional validation across three distinct murine models of DSS-induced colitis of increasing biological complexity: the original gnotobiotic defined microbial community (**Figure 4A)**, specific-pathogen-free mice with a native microbiota (**Figure 4D**), and an extended DSS recovery model (**Figure 5A**).

**Figure 4.**
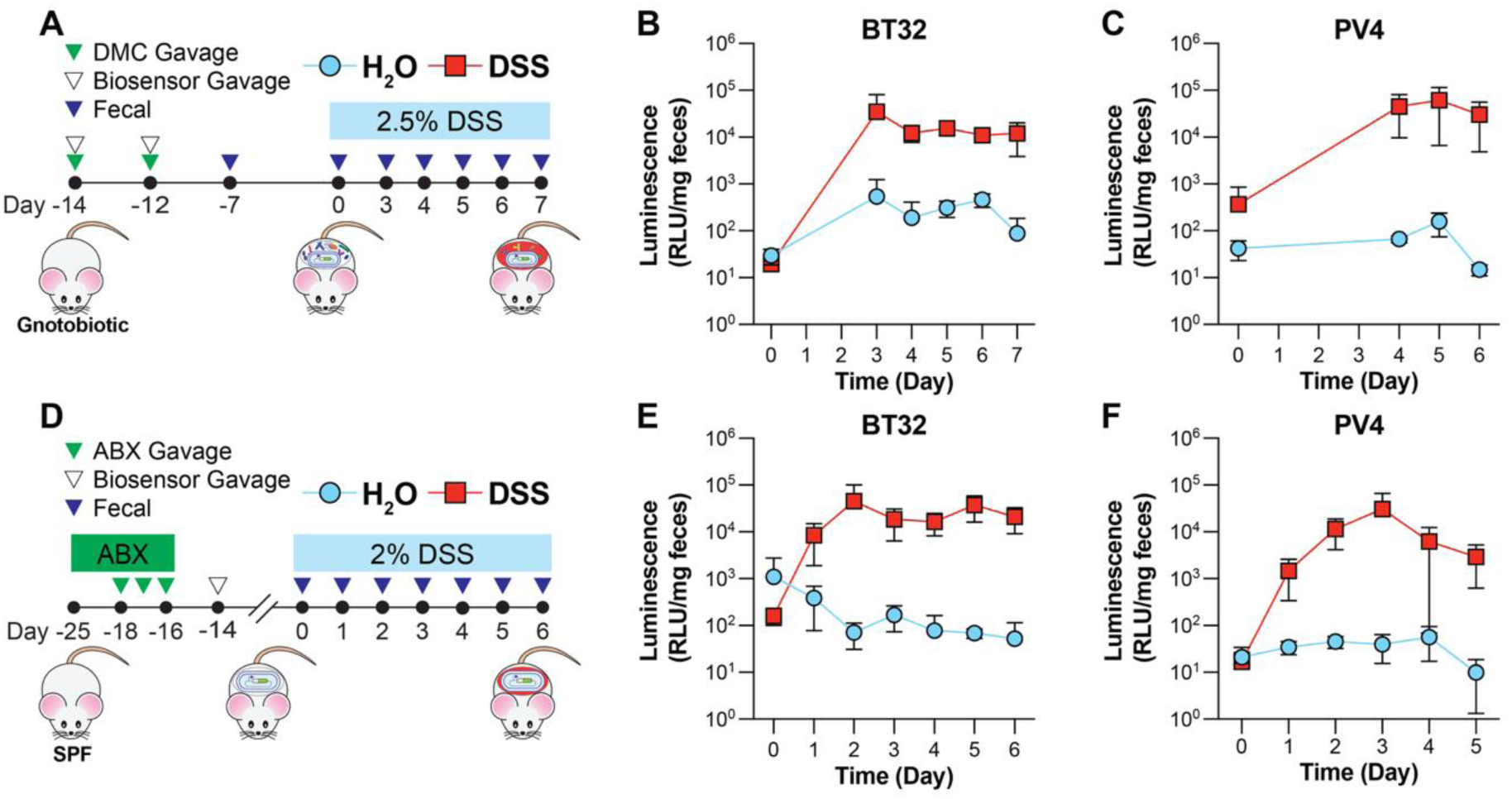
BT32 and PV4 function as biosensors for DSS-induced colitis in gnotobiotic and conventional mice. (**A**) Experimental timeline for validation of BT32 and PV4 biosensors in a gnotobiotic DSS model. Biosensor (BT32 or PV4) was gavaged alongside the remaining DMC members twice, at least two days apart and at least two weeks before DSS administration. Engraftment of biosensor plus all DMC members was checked one week after initial gavage. Mice were administered 2.5% DSS or regular drinking water for up to 7 days. Fecal samples were collected each day starting on Day 3. (**B**) Luminescence (RLU per mg of collected feces) of BT32 in H_2_O (n = 3) and DSS (n = 4) over time. (**C)** Luminescence (RLU per mg of collected feces) of PV4 in H_2_O (n = 3) and DSS (n = 3) over time. **(D**) Experimental timeline for validation of BT32 and PV4 biosensors in SPF DSS Model. Mice are treated with antibiotic water (ciprofloxacin-HCl, metronidazole) (ABX) for one week, with daily gavages of antibiotics (metronidazole) for 3 days at the end of treatment. After a 48-hour washout, biosensor (BT32 or PV4) was gavaged at least two weeks prior to DSS administration. Mice were administered 2% DSS or regular drinking water for up to 6 days. (**E**) Luminescence (RLU per mg of collected feces) of BT32 in H_2_O (n = 3) and DSS (n = 5) over time. (**F)** Luminescence (RLU per mg of collected feces) of PV4 in H_2_O (n = 4) and DSS (n = 4) over time.

**Figure 5.**
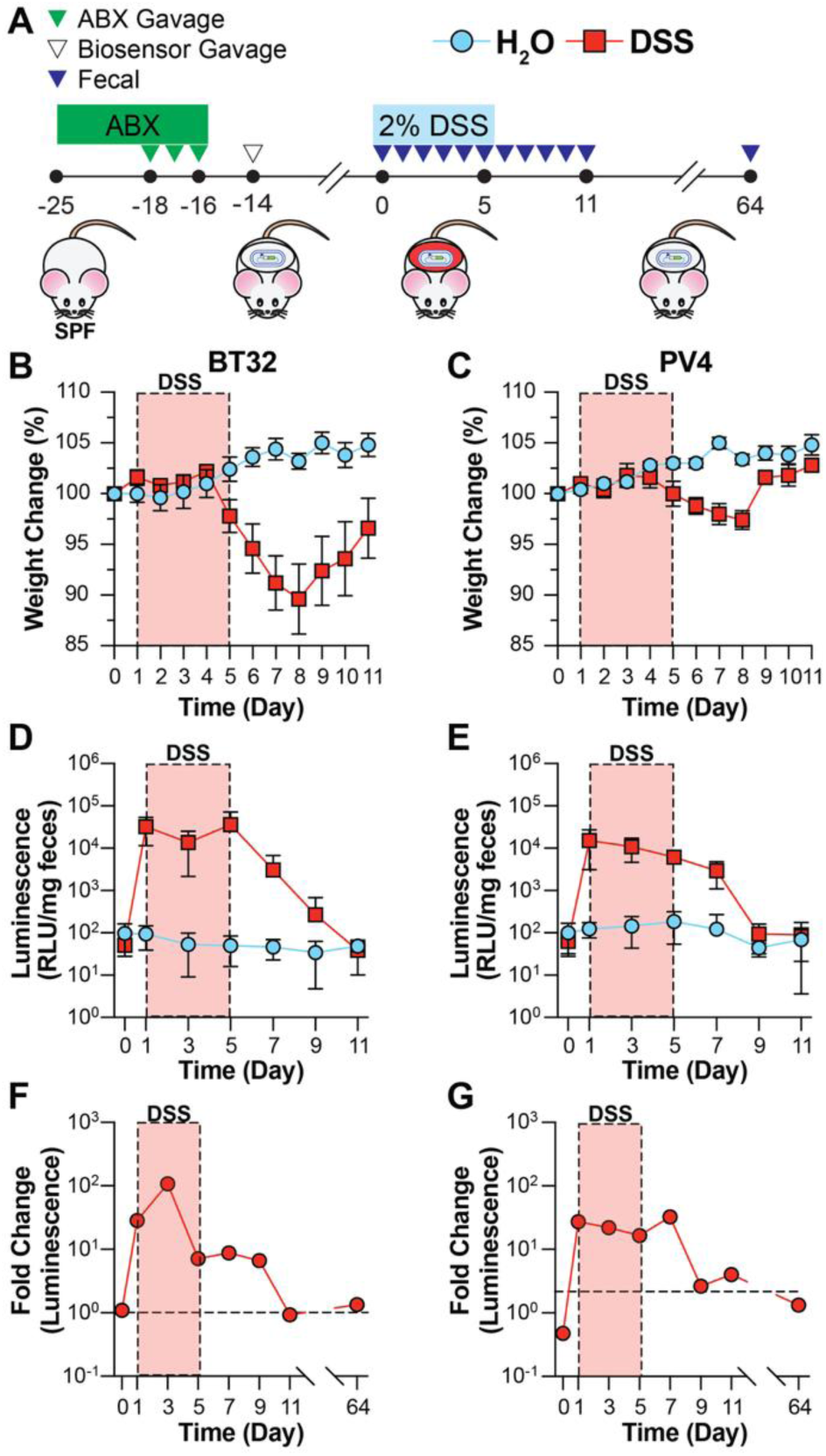
BT32 and PV4 biosensors respond to inflammation and return to baseline in DSS recovery model. (**A**) Experimental timeline for validation of BT32 and PV4 biosensors in a DSS recovery model. Conventional mice were treated with antibiotic water (ciprofloxacin-HCl, metronidazole) (ABX) for one week, with daily gavages of antibiotics (metronidazole) for 3 days at the end of treatment. After a 48-hour washout, biosensors (BT32 or PV4) were gavaged at least two weeks prior to DSS administration. Mice were administered 2% DSS or regular drinking water for 5 days, then returned to regular drinking water to recover. Mice were maintained until Day 64 post DSS administration to ensure long term biosensor engraftment. (**B**) BT32 and (**C**) PV4 DSS groups lose weight over time compared to H_2_O control groups up to Day 8, as a measure of percent change in body weight relative to baseline (Day 0), then begin to recover and gain weight to Day 11 (n = 5 per group). (**D**) BT32 and (**E**) PV4 biosensors response to DSS conditions and recovery as measured by Luminescence (RLU per mg of collected feces). BT32 and PV4 sustain response post return to regular drinking water. (**F**) BT32 and (**G**) PV4 fold change in DSS conditions calculated by RLU per CFU, to control for biosensor abundance in fecal contents. Elevated fold change is sustained after return to regular drinking water, and the return to baseline corresponds to weight recovery. Both BT32 and PV4 biosensors are at or below baseline by Day 64.

In the gnotobiotic DMC model, germ free mice were stably colonized with the engineered Bt-BT32 and Pv-PV4 biosensors alongside the other 10 DMC members two weeks prior to 2.5% DSS administration (**Figure 4A**). DSS-treated cohorts developed characteristic disease phenotypes, including weight loss, bloody stool, and colon shortening (**Figure S16)**. Both biosensors were tracked by measuring fecal luminescence each day, and maintained low baseline expression prior to DSS administration, followed by rapid and robust signal amplification by Day 3-4 (**Figure 4B-C).** After normalization to biosensor abundance, the sensors achieved a 100-fold (BT32) and 75-fold (PV4) induction over healthy controls (**Figure S16 C-D, G-H)**. Throughout the duration of the study, untreated cohorts maintained a low baseline luminescence. Next, to evaluate each biosensor’s utility in an undefined microbial environment, each was engrafted into conventional SPF mice following an antibiotic pretreatment regimen to deplete endogenous Bacteroidales spp. and enable engineered strain engraftment. Robust biosensor activation was observed 24 hours after 2% DSS administration and remained elevated throughout treatment (**Figure 4E-F**), approximately 100-fold over controls with some measurements exceeding 1,000-fold (**Figure S17**). Together, the BT32 and PV4 biosensors rapidly respond to the onset of DSS treatment in both simplified and complex microbial communities.

A critical feature of a function biosensor is its capacity to return to a baseline “off” state following disease resolution. To evaluate these kinetics, engrafted SPF mice were administered 2% DSS for 5 days and subsequently returned to standard drinking water to initiate recovery. Both biosensors responded within 24 hours, and the signal remained elevated until mice began regaining weight around Day 8 (**Figure 5B-C**). The BT32 reporter successfully returned to true baseline by Day 11, while the PV4 signal remained slightly elevated compared to untreated controls, but much lower than during disease (**Figure 5D-G)**. Notably, at Day 64 post-DSS treatment, both engineered biosensors remained stably colonized and expression in formerly DSS-treated mice was indistinguishable from untreated controls (**Figure 5F-G, Figure S18).** Collectively, across all three models, Bt and Pv engineered with the ECF-type sigma factor operon promoter demonstrated high diagnostic specificity, sensitivity, stable long-term colonization, and return to baseline following disease resolution.

## Discussion

Mining microbial transcriptomes *in vivo* is a powerful and generalizable approach for the discovery of functional biosensors of complex gut states. By sampling host transcriptomes during health and disease, we identified candidate sensor systems (CSSs) that might be overlooked by traditional, single analyte *in vitro* discovery screens. This strategy uses natural bacterial responses, along with operon structure and regulatory context, to prioritize candidate sensors most likely to respond to *in vivo* perturbations. Using our pooled BCSeq screening platform, we systematically profiled almost 200 endogenous promoters across two gut-resident

Bacteroidaceae chassis in *in vitro*, healthy, and disease conditions and identified 8 CSSs that respond to canonical physiological changes associated with gut inflammation, including oxidative stress, acidic pH, and altered bile acid exposure. However, the strongest candidates, BT32 and PV4, illustrate the value of this approach, as both are robustly induced during intestinal inflammation, yet do not respond to canonical inflammation stressors *in vitro.* Individual *in vivo* validation across gnotobiotic, SPF, and disease recovery murine models demonstrated that the BT32/PV4 biosensors provide rapid activation, high diagnostic sensitivity, stable colonization, and a return to baseline expression after disease recovery. These findings underscore that gut-resident symbionts harbor a reservoir of clinically-relevant sensor systems that can be unlocked by mining natural transcriptional responses to complex disease environments and converted into engineered sensors of intestinal physiology. Importantly, this approach is not limited to this specific disease model or bacterial chassis and can be adapted to fit other perturbations, disease contexts, or microbial taxa of interest.

Our *in vitro* culture assays and genetic deletions established a regulatory connection between the *BT_4647-49* and *BVU_2338-40* operons and endogenous SPT-mediated sphingolipid metabolism. Because these operons fail to respond to canonical membrane stressors, including oxidative stress, acidic media, heat, or bile acids, we posit their activation is likely driven by a membrane state or disruptions tied to sphingolipid flux, rather than a general periplasmic protein misfolding. This three-gene operon architecture is highly conserved within the Bacteroidales, an order well known for *de novo* sphingolipid synthesis in the human gut microbiome^78,83^. *Bacteroides*-derived sphingolipids have established roles in gut homeostasis, immune modulation, and outer membrane vesicle (OMV) signaling^79,82,84^. Further, both host and bacterial sphingolipids and ceramides are upregulated in patients with inflammatory bowel disease^6,85–87^, suggesting that BT32/PV4 activation is relevant to gut inflammation. Furthermore, published transcriptomic data from anti-CD3 and flagellin-induced immune activation models using a defined microbial community demonstrate the upregulation of a homologous operon in *Bacteroides sartorii*^4^, suggesting a functional role for this operon across diverse *in vivo* inflammatory environments. Notably, BT32/PV4 activation preceded inflammatory phenotypes *in vivo*, indicating that these systems respond to early manifestations of epithelial disruption rather than downstream inflammation byproducts like nitrate^20,24,39^, thiosulfate, or tetrathionate^21,22^. Although further mechanistic investigation is warranted, these biosensors could thus be potential indicators of epithelial disruption itself, which would be useful in diagnostics, recovery monitoring, therapeutic assessment, and sense-and-respond live biotherapeutics.

Despite our investigation into BT32/PV4 activation mechanism, the full dynamic range of the *in vivo* response could not be recapitulated *in vitro*. Chemical and genetic inhibition of sphingolipid biosynthesis modulated expression, but the resulting fold changes did not approach *in vivo* levels, indicating that additional host- or microbiome-derived physiochemical factors contribute to full activation. This gap reflects the broader premise motivating our approach that the gastrointestinal tract is a biochemically complex ecosystem whose native signals are difficult to reconstruct outside of the host. More broadly, *in vivo* bacterial transcriptomics is inherently complex, subject to community composition effects, engraftment variability, and condition specific noise. By pairing transcriptomic profiling with high-throughput *in vivo* functional screening, we demonstrate an efficient route to biosensor discovery that can capture the breadth of biomolecular changes in the gut due to disease. Additionally, we show that non-model gut symbionts harbor sensitive, sophisticated regulatory circuits tailored to complex disease states. This work establishes *in vivo* transcriptomic mining as a generalizable strategy for discovering biosensors in gut-resident bacteria and provides a scalable foundation for engineering the next generation of autonomous, commensal-based diagnostics and smart therapeutics for intestinal disease.

### Limitations and Future Directions

Although this pipeline successfully identified highly responsive, non-model biosensors, several technical and biological limitations remain. Technically, barcode screening *in vivo* required substantial amounts of RNA and reagents to process CSS libraries without over amplification. Application of this method to lower-biomass environments or low abundance community members requires further optimization. Despite our efforts to identify the inducer signal and potential links to sphingolipid biosynthesis, we were unable to identify a single analyte that activated the validated ECF regulatory system, limiting our understanding of their full dynamic range. With a known defined analyte, future work could incorporate promoter engineering, directed evolution^88,89^, and AI-guided design^90^ to expand the biosensor’s dynamic range. Future work could also expand to include the mining and screening of downregulated genes, using a repressor-based circuit for screening^38^. Finally, while we focused on the dominant order Bacteroidales, prominent taxa more suited to other disease models, including gut Clostridia and Lachnospiraceae, could be mined and engineered using this approach in tandem with newly developed genetic engineering techniques^91,92^.

Additionally, the DSS colitis model itself also carries limitations to translational applications. Engineered Bacteroidales strains achieved stable long-term engraftment in conventional SPF mice, but only after significant antibiotic conditioning prior to biosensor introduction. Clinical translation will require either improved or engineered engraftment strategies or biosensors capable of transient sensing without persistent colonization. The biological heterogeneity of the gastrointestinal tract also presents distinct challenges for living diagnostics. Intestinal microenvironments vary drastically along both the longitudinal and radial axes and localized cellular behavior may introduce spatial signal variance. Finally, DSS-induced colitis, while a useful model for engineering-cycle discovery, does not fully capture the relapsing-remitting etiology of human IBD, and the resulting biosensor responses may not generalize to other inflammatory contexts. Applying this transcriptomic mining approach to genetic, infectious, and dietary models of intestinal disease will be important to establish the broader generalizability of the method.

## Acknowledgements

We would like to thank the following people who helped contribute to this paper: Bana Jabri for the original germ-free breeding pairs; Eric Pamer, Sam Light, Laurie Comstock, Tatyana Golovkina, and Alexander Chervonsky for providing germ-free breeding pairs and mice; Becky Ortiz, Emma Elmiger, Yessenia Sierra, Catherine Hoyle and Kristin Kolar at University of Chicago Gnotobiotic Research Animal Facility; Anitha Sundararajan, Huaiying (Eddi) Lin and Ramanujam Ramaswamy at University of Chicago DFI Microbiome Metagenomics Facility for their help with RNA-seq analysis; Christian Jacoby and Sam Light for help with APOL2 protein; Jack Arnold for the Prokka mapped genomes; Rory McGann for help preparing the DMC consortiums.

The authors are supported by the National Institute of General Medical Sciences of the National Institutes of Health (R35GM147478; M.M.), by the National Institutes of Health (IMSD Program 5R25GM109439-07 and Molecular and Cellular Biology training program T32 GM007183 to J.F.-S.), by the Arnold and Mabel Beckman Foundation through the Beckman Young Investigator Program (M.M.).

## Author Contributions

J.G., D.V., and M.M. designed and performed experiments. J.G., S.M., J.G., J.F-S. and M.M. performed mouse experiments. J.G. performed data analysis. J.G. and M.M. conceived this study, analyzed the data, discussed results, and wrote the manuscript.

## Declaration of generative AI and AI-assisted technologies in the writing process

During the preparation of this work, the authors used Claude (Anthropic) to improve the readability of the manuscript. After using this tool/service, the authors reviewed and edited the content as needed and take full responsibility for the content of the published article.

## STAR Methods

### RESOURCE AVAILABILITY

#### Lead Contact

Further information and requests for materials and reagents can be directed to lead contact Mark Mimee.

#### Materials Availability

Bacterial strains and genetic constructs are available upon request.

#### Data and Code Availability

Code is available upon request. Genbank accession numbers for bacterial genomes are included in **Table S1**. RNAseq (GSE343114) and BCseq (GSE343117) data are deposited in GEO.

### METHODS

#### Bacterial Strains and Growth Conditions

All strains, and their respective medias, used are listed in Table S1. Anaerobic strains were grown anaerobically at 37°C in indicated media in a Coy anaerobic chamber with an atmosphere of 85% N_2_, 5% H_2_, 10% CO_2_. Plates and media were pre-reduced overnight in anaerobic atmosphere before inoculation, plating, or restreaking of cultures. The following media were used: *BHIS Media:* 37g Bacto BHI Premix, 5g Yeast extract (Gibco), 4ml Resazurin (Arcos Organics) at 250 µg/mL, and 10 mL L-cysteine-HCl (EMD Millipore) at 50 µg/mL, 1 mL menadione (MP Biomedical) at 1 mg/mL, 10 mL Hemin (Sigma Aldrich) at 1 mg/mL, and 20 mL Trace Mineral Supplements (ATCC) were added after autoclaving. BHIS plates were made with the same recipe, substituting BHI Premix for BHIS Agar Premix. *BHI Media:* 37g Bacto BHI Premix, and 1 mL menadione (MP Biomedical) at 1 mg/mL and 10 mL Hemin (Sigma Aldrich) at 1 mg/mL were added after autoclaving. *Bacteroides Minimal Media (B-MM):* Ammonium sulfate (1 g/L), Sodium carbonate (1 g/L), Potassium phosphate buffer (pH 7.3, 80 mL/L), Vitamin B12 (0.5 mL/L), Iron sulfate heptahydrate (0.4 mg/mL, 1 mL/L). 10 mL Glucose (1M), 10 mL L-cysteine-HCl (EMD Millipore) at 50 µg/mL, 1 mL menadione (MP Biomedical) at 1 mg/mL, 10 mL Hemin (Sigma Aldrich) at 1 mg/mL, and 20 mL Trace Mineral Supplements (ATCC) were added after autoclaving. *MRS and Reinforced Clostridial Media (RCM):* Prepared from premixed powders (Thermo Fisher). 10 mL L-cysteine-HCl (EMD Millipore) at 50 µg/mL added after autoclaving.

Unless otherwise noted, *E. coli* strains were grown aerobically at 37°C in LB broth while shaking at 250 RPM (BD Difco) or on plates (agar 15 g/L) (Fisher Scientific). *E. coli* S17 strains were grown on LB + carbenicillin (Cb) (100 µg/mL) plates and in LB + Cb media for selection during cloning and conjugation processes. Where indicated, antibiotics were added to the growth medium or plates to the following final concentrations: carbenicillin (Cb, 100 µg/mL), erythromycin (Em, 25 μg/mL), gentamicin (Gm, 200 µg/mL), anhydrous tetracycline (aTc, 100ng/mL).

#### Defined Community Preparation

The Defined Microbial Community (DMC) is a consortium of 13 species, listed in Table S1. The DMC was constructed by growing each microbial member 24-48 hours (anaerobically for all but *E. coli)*. Most anaerobic bacteria members were grown in BHIS media. For *A. muciniphila*, N-acetyl-glucosamine (Sigma) was added to 10 mM to BHIS. *For L. reuteri*, MRS (Oxoid) plates and liquid were made from premixed powder. For *B. adolescentis*, Reinforced Clostridial Media (RCM; BD) was used for plates and MRS used for liquid culture growth. *E. coli* was grown aerobically in LB media broth or on plates at 37°C with 250 rpm shaking. To prepare the DMC for engraftment into gnotobiotic mice, each culture was normalized to the same background-adjusted optical density and equal volumes of each culture were combined together. The pooled consortia was stored at −80°C by combining 750 µl of cells with 250 µl of 50% glycerol.

#### Amplification of Genetic Parts and Cloning

PCR amplification was performed using two polymerases depending on the template source. Genetic parts were amplified from plasmid templates using Q5 High-Fidelity DNA Polymerase (New England Biolabs) according to the manufacturer’s protocol. Reactions were prepared in a total volume of 50 µL containing 1× Q5 reaction buffer, 200 µM dNTPs, 0.5 µM of each primer, 1 ng template DNA, and 1 U Q5 polymerase. Cycling conditions consisted of an initial denaturation at 98°C for 30 seconds, followed by 30-35 cycles of denaturation at 98°C for 10 seconds, annealing at 60°C for 30 seconds, and extension at 72°C for 30 seconds per kb, with a final extension at 72°C for 2 minutes. Genetic parts amplified from *Bacteroides thetaiotaomicron* and *Phocaeicola vulgatus* genomic DNA were amplified using KAPA Robust DNA Polymerase (Roche) according to the manufacturer’s protocol. Reactions were prepared in a total volume of 50 µL containing 1× KAPA Robust buffer, 0.3 µM of each primer, and 10-100 ng template DNA. Cycling conditions consisted of an initial denaturation at 95°C for 3 minutes, followed by 35 cycles of denaturation at 95°C for 15 seconds, annealing at 60°C for 15 seconds, and extension at 72°C for 15 seconds per kb, with a final extension at 72°C for 1 minute. All PCR products were verified by agarose gel electrophoresis prior to downstream use.

#### Bacteroidales Strain Engineering

Genetic parts including primers, plasmids, promoters, and homology regions for deletion mutants are in the Supplementary Tables S2-S5. Plasmids assembled by Gibson or Golden Gate assembly were transformed into electrocompetent *E. coli* S17 using electroporation, recovered for 30-60 minutes in LB, and plated on LB+Cb plates. Colonies were verified for successful transformation via Colony PCR and plasmids were purified via Qiaprep Spin Miniprep Kit (Qiagen) for validation with Sanger sequencing. Validated colonies were grown overnight for conjugation with Bacteroidales species. Recipient Bacteroidales (Bt or Pv) and donor *E. coli* S17 were grown overnight in respective media. 250 µL of *E. coli* were spun down for 5 min at 5,000xg, supernatant discarded, and then resuspended in 1 mL of Bacteroidales. The combined cultures were spun down, supernatant discarded, and resuspended in 25 µL of BHIS media for spot plating on BHIS plates (no antibiotics). After drying, mating plates were incubated aerobically overnight. Following incubation, mating spots were scraped using sterile inoculation loops into 1 mL of BHIS and resuspended. 200 µL of the suspension was plated on BHIS Em/Gm plates and grown anaerobically for 48 hours. Single colonies were selected, restreaked for isolation on BHIS Em/Gm plates and incubated for 48 hours. Colonies were confirmed via luciferase measurement, colony PCR, or Sanger sequencing on isolated gDNA. All strain engineering was done via these methods, except for the Oligopool Pool2 and Pool3 library prep.

#### Bacteroides Promoter Libraries

For Oligopool library cloning, *E. coli* S17 were transformed with the CSS library, inserted into the pNBU1-CSS-NL backbone via Golden Gate assembly. Transformations were recovered for 1 hour in LB and then plated on 8-10 LB + Cb plates at a dilution that yielded individual colonies (∼500-1000 per plate). Plates were scraped, resuspended in Lb+Cb media, labeled as ‘Transformation Starters’, and saved at −80°C with an equal volume of 50% glycerol added. Transformation Starters were thawed, spun down to remove glycerol, diluted 1:20 in LB + Cb media, and grown to mid-log phase before mating with overnight Bt or Pv as described above. Multiple spots were seeded, incubated, and scraped per library preparation. After scraping, cells were thoroughly resuspended, diluted, and plated on 8-10 BHIS Em/Gm plates to yield individual colonies (∼500-1000 per plate). After 24-48 hours of growth, 3 mL of BHIS + Em/Gm media was added and each plate scraped to collect all transformed colonies. Cells were resuspended thoroughly in BHIS + Em/Gm media, labeled as Bt and Pv Library Starters, and saved at −80°C with an equal volume of 50% glycerol added. To prepare libraries for gavage, 2mL stocks were thawed, diluted 1 to 20 in BHIS Em/Gm media and grown overnight. For *in vitro* analysis, overnight library cultures prepared for *in vivo* gavage were subcultured 1:50 and grown to mid-log for RNA and gDNA isolation and BCSeq analysis.

#### Knockout Mutants

Homology arms for Deletion Mutants (Table S5) 1 kb upstream and downstream of the target deletion, were amplified from host gDNA and inserted into pLGB13^54^ using Gibson Assembly. Successfully conjugated Bt and Pv colonies were restreaked onto BHIS + aTc (100 ng/µL) plates to select deletion mutants. Colonies grown on BHIS + aTc plates were screened via colony PCR for deletion of the target genes and restreaked on BHIS plates for isolation.

#### Animal Procedures

All animal experiments were carried out in compliance with the University of Chicago Institutional Animal Care and Use Committee (IACUC) (Protocol 72610) guidelines and maintained at the University of Chicago Animal Resource Center (ARC). All experiments were carried out in C57BL/6J mice. Germ-free and gnotobiotic mice with a defined consortium of microbes^46^ bred and housed within the University of Chicago Gnotobiotic Research Animal Facility (GRAF). Specific pathogen-free (SPF) mice were purchased from the Jackson Laboratory and maintained in the University of Chicago SPF facility.

All experiments were performed in accordance with the Guide for the Care and Use of Laboratory Animals (8th ed.) and were approved by the Institutional Animal Care and Use Committee of the University of Chicago (Protocol 72610).

#### Gnotobiotic Defined Microbial Community (DMC)

Gnotobiotic mice associated with a stable consortium of 13 microbes were bred and housed within the University of Chicago Gnotobiotic Research Animal Facility. Breeding pairs of mice were gavaged with 200 µl of thawed DMC twice, separated by three days. The presence of each bacteria in the gut was determined by 16S rRNA sequencing of fecal pellets. Community composition was vertically transmitted to pups and remained stable for >3 years, as monitored by 16S rRNA sequencing of fecal samples with the aid of the UChicago Duchoissois Family Institute Microbiome Metagenomics Facility (DFI MMF). During this process, the bacterium *Turicibacter sanguinis* was acquired. Gnotobiotic C57BL/6 mice were bred and housed in Trexler-style flexible firm isolators (Class Biologically Clean) within Ancare polycarbonate mouse cages (N10HT). Gnotobiotic mice were weaned at 21 days of age and were fed an autoclaved, plant-based mouse chow (LabDiet JL Rat and Mouse/Auto 6F 5K67). All mice were housed in cages containing autoclaved Teklad Pine Shavings (cat# 7088) with a 12-hour light/dark cycle at a standard room temperature of 20-24°C. All mice were euthanized by CO_2_ asphyxiation followed by cervical dislocation as a secondary measure. Experiments utilized both male and female mice, except during individual validation models (male only). This gnotobiotic DMC model was used for DSS transcriptomics. For *in vivo* vs *in vitro* transcriptomics, two additional bacteria, *Bacteroides fragilis* and *Coporrococcus comes,* were engrafted before sampling 14 days later.

#### Gnotobiotic DSS Model

8–12-week-old C57BL/6 mice colonized with the 13-member DMC in the UChicago Gnotobiotic Research Animal Facility were given 2.5% Dextran Sodium Sulfate (DSS) drinking water and monitored for 6-7 days. DSS drinking water was prepared by adding dextran sulfate sodium salt (M.W. approx. 36000 - 50000, Colitis Grade, Fisher Scientific) in deionized water and filter sterilized prior to administration to mice. Each day, mice were monitored for weight, decline in body conditions, lack of grooming, dehydration, and lethargy. At 2.5% DSS, mild to moderate bleeding in fecal samples was observed by Day 5-6. Animals were removed from study if the following endpoints occurred before Day 6 for males or Day 7 for females: 15% loss in body weight, rectal prolapse, severe decline in body conditions/activity, overt bleeding. On Day 6 (males) or Day 7 (females) mice were euthanized to measure colon length and intestinal contents collected for analysis.

#### CSS Engineered Pool Screening

DSS Candidate Libraries were screened by gavaging 6–8-week-old C57BL/6 germ-free mice with 100 µL of pooled Bt and Pv candidates, alongside 100 µL of the remaining community members. Gavages were repeated 2-3 days later. Fecal pellets were collected between Day 3 and Day 7 to confirm engraftment. Generally, additional procedures were not performed in the first to week post-gavage to allow the community to reach equilibrium. Two to three weeks following gavage, mice administered 2.5% DSS drinking water and monitored for symptoms.

#### Specific Pathogen-free (SPF) DSS Model

5–6-week-old C57BL/6 mice were purchased from Jackson Laboratories, allowed to acclimate for at least 3 days and provided antibiotic-supplemented water (ciprofloxacin-HCl (0.625 g/L), metronidazole (1 g/L), and 3% sucrose in reverse osmosis water, filter sterilized) for 7 days. On Days 8-10, drinking water was supplemented with only ciprofloxacin-HCl (0.625 g/L) and mice were gavaged daily with metronidazole (100 mg/kg). After a 48-hour washout period, mice were gavaged with 200 µL of overnight grown Bt or Pv cultures. Engraftment of bacteria was confirmed one week after gavage by selective plating on BHIS + Em/Gm. Two weeks after gavage, drinking water of select cages was supplemented 2% DSS. All mice were weighed daily and monitored for symptoms. Endpoints were the same as the gnotobiotic DSS model. To measure fecal luminescence and engineered strain burden, fresh fecal pellets were weighed, homogenized in 200 µL PBS, and spun down (5,000 x g for 30 seconds) to collect supernatant for RLU measurement and CFU plating. Mice were sacrificed on Day 6 (male) or Day 7 (female) of DSS treatment.

#### DSS Recovery Model

5–6-week-old SPF C57BL/6 mice were ordered from Jackson Laboratories, treated with antibiotic supplemented water and gavaged with biosensor strains as described above. Two weeks after gavage, drinking water of select cages was supplemented with 2 % DSS for 5 days and then returned to regular drinking water for the remainder of the experiment. All mice were weighed daily and monitored for symptoms. Fecal pellets were collected every other day to measure luminescence and Bt/Pv burden for 11 days. After Day 11, mice were maintained for 50+ more days to confirm long term engraftment and sensor return to baseline.

#### Isolation and analysis of Mouse contents

In all models, when indicated, cecal and colon contents were weighed, resuspended in 200-500 µL of PBS and homogenized in a PowerLyzer 24 at 2,000 RPM for 2 minutes. Homogenized fecal pellets were spun down for 30 seconds at 5,000 x g and the supernatant used for relative luminescence, lipocalin-2 ELISA, and colony forming unit (CFU) measurements.

#### 16S rRNA Sequencing

DNA extraction from fecal samples was performed using the QIAamp PowerFecal Pro DNA Kit (Qiagen) and the V4-V5 region of 16S rRNA-genes were PCR amplified using barcoded dual-index primers. Illumina compatible libraries were generated using the Qiagen QIASeq 1-step amplicon kit. Library quality control was performed using Qubit and Tapestation and sequenced on the Illumina MiSeq platform in the Duchossois Family Institute Microbial Metagenomics Facility using a 2×250 Paired End reads, generating 5,000–10,000 reads per sample. Raw V4-V5 16S rRNA gene sequence data was demultiplexed and processed through the dada2 (v1.18.0)^93^. pipeline into Amplicon Sequence Variants (ASVs).

ASVs were identified with the RDP classifier (v2.13)^94^ up to the genus level with a minimum bootstrap confidence score of 80. ASVs were subsequently BLASTed against a custom database of 16S rRNA sequences corresponding to the members of the defined microbial community and assigned to specific species.

#### Genomic DNA Isolation

Bacterial Genomic DNA (gDNA) was isolated from *in vitro* cultures using the Qiagen Powerlyzer Kit. Bacterial gDNA was isolated from mouse fecal samples using the Qiagen PowerMicrobiome Kit. DSS samples were cleaned up a second time using Zymo DNA Clean and Concentrator to remove remaining DSS.

#### RNA Isolation

All steps involving RNA handling were performed with sterile filter pipette tips (Fisher Scientific) and RNase Away spray was used to clean surfaces.

##### *In vitro* Bacterial Cultures

Overnight cultures were diluted 1:50 in BHIS media and grown to mid-log phase (0.4 – 0.7 OD_600_). RNAProtect (Qiagen) was added to the cells per manufacturers protocols prior to RNA isolation. Total RNA was isolated and purified using RNAeasy RNA extraction kit (Qiagen).

##### Cecal Contents

Mouse cecal contents were processed immediately (less than 10 minutes from euthanasia) and extracted by acidic phenol: chloroform extraction (Phenol:Chlorofrom:IAA 125:24:1 pH 4.5, Ambion 9720), followed by a Qiagen RNeasy column purification. DSS samples were precipitated with lithium chloride to remove DSS prior to DNAse treatment. Isolated RNA was resuspended in lithium chloride to a final concentration of 2.5M and incubated for at least 30 minutes at −20°C (preferably overnight). After incubation, precipitated RNA was pelleted and washed with 70% ethanol twice, air dried, and resuspended in nuclease-free water. All isolated RNA was DNase treated (TURBO), column purified (Zymo), and concentration was determined by NanoDrop before further analysis.

#### RNA sequencing

RNA was submitted to the UChicago DFI Metagenomics Core for Illumina RNA-seq Library Prep. Illumina compatible libraries were generated using NEBNext® Ultra™ II Directional RNA Library Prep Kit. Ribosomal RNA depletion (NEB) was used prior to library preparation. Libraries were sequenced using 2×150bp reads on NextSeq 1000 at the Duchossois Family Institute Microbiome Metagenomics Facility or equivalent at SeqCenter.

#### Barcode Sequencing

BCSeq methods were adapted from previously published methods^31,95^. 25-30 μg of RNA was DNAse treated with TURBO as follows: 15 µL buffer, 6 µL TURBO, RNA + nuclease-free water H_2_O to 150 µL, incubated at 37°C for 30-60 minutes. DNAse-treated RNA was purified using Zymo RNA Clean and Concentrated 25 Kit and eluted in 50 µL. Next, DNase-treated RNA was quantified via Nanodrop and up to 5 μg was used for targeted reverse transcription (D105, Superscript IV) at the following concentrations: 10 µL DNase treated RNA, 1 µL H2O, 1 µl 10 μM dNTP, 1 µL D105 (20 µM). The mixture was incubated for 5 min at 65°C and then for 1 minute on ice. Next, a SSIV Master Mix (4 µL Buffer, 1 µL DTT, 1 µL SSIV, 1 µL RiboLock per reaction) was added and samples incubated at 55°C for 90 min, followed by a 15 min inactivation step at 80°C. 1 µL of RNase H was added and incubated for another 30 minutes at 37°C. For each sample, 4 separate reverse transcription reactions were performed, pooled, and then purified using Zymo DNA Clean and Concentrator with a 20 µL elution volume. To ligate adapter sequences, 5 µL cDNA and 2 µL adapter (40 µM) were incubated at 75°C for 3 min then put on ice. T4 RNA Ligase Master Mix (2 µL Buffer, 0.8 µL DMSO or H_2_O, 0.2 µL ATP (100 mM), 1.5 µL Ligase, and 8.5 µL PEG 8000 (50%) per sample) was added and incubated overnight at 22°C. 4 ligations were performed per sample, pooled, and purified using Zymo columns. Libraries were prepared by first running qPCR (SYBR, Quantstudio 3) to determine the number of cycles before exponential amplification (about Ct+1 in most cases) and to confirm proper library size (melt curve, single peak). Libraries were then PCR amplified with partial Illumina adaptors (Q5), pooled (6 reactions each), purified (Zymo DNA columns), and quantified via Qubit. Submission to Azenta/Genewiz for EZ Amplicon sequencing required 20 µL volume of at least 500 ng library DNA.

#### Relative Luminescence (RLU)

Equal volumes of cell culture or resuspended fecal/cecal contents supernatant and Nano-Glo working reagent (1:50; Nano-Glo Substrate: Nano-Glo buffer) (Promega) were added to a flat white 96 well plate (Costar) and luminescence was recorded (Tecan infinite 200). Normalized Relative Luminescence for *in vitro* cultures was calculated by subtracting background RLU and OD_600_ from measured values and then dividing RLU by OD_600_. To calculate *in vitro* fold change, *in vitro* normalized luminescence for the induced condition was divided by the uninduced or negative control condition. Normalized Relative Luminescence for *in vivo* samples was calculated by subtracting background RLU from measured values and then dividing RLU by feces mass. To calculate *in vivo* fold change, *in vivo* luminescence was normalized to bacterial load by dividing the background-subtracted RLU by the corresponding CFU (RLU/CFU) and then averaging DSS RLU/CFU divided by Healthy RLU/CFU.

#### *In vitro* Induction Assays

Selected CSS were evaluated for potential inducers by either stimulating mid-log growth cultures with an inducer or by subculturing overnight cultures in the presence of the inducer. For mid-log induction, overnight cultures of CSS are subcultured 1:50 in respective media and grown to mid-log phase (0.4 - 0.8 OD in BHIS, 0.4 - 0.6 in B-MM). At mid-log, the following concentrations of inducers were added: TPEN (10 µM), APOL2 (10 µg/mL), Bile Acids (0.03% w/v), Polymyxin B (0-500 µg/mL), and Myriocin (0-50 µM). Induction with oxygen was done by splitting aerobically grown cultures and shaking one half aerobically at 250 rpm. For Biolog plates, cultures were grown overnight in provided Biolog media, subcultured until mid-log growth, then added to the selected plates. For each induction assay, RLU and OD_600_ were measured two hours after induction. For subcultures in the presence of potential inducers, overnight cultures were diluted 1:20 in media supplemented with inducers. CSS are measured 4, 8 and 20-24 (overnight) hours post induction. For all assays, cultures with no inducer or equal volumes of inducer solvent (i.e. PBS, DMSO, Media) were included to calculate fold change. Fold Changes were calculated by comparing normalized relative luminescence from induced to uninduced cultures. For Biolog plates, fold change was calculated by comparing individual wells normalized luminescence to the plate median. When indicated for all experiments, fold change was log_2_ transformed for plotting purposes and minimum thresholds for induction were set at Log_2_Fold Change > 1.

#### Lipocalin-2 ELISA Assay

Supernatant from homogenized mouse fecal, cecal, or colon contents was diluted 1:100 or 1:500 in PBS for lipocalin-2 measurement via sandwich ELISA using the Mouse Lipocalin-2/NGAL DuoSet ELISA kit and manufacturer’s instructions (R&D Systems).

#### Quantitative PCR (qPCR)

Each qPCR reaction contained 1-2 µL cDNA library, 1 µL each primer (10-20 uM), 10 µL Power UP SYBR Green Master Mix, and molecular grade water to 20 µL total. Reactions were run on a QuantStudio 3 System (Applied Biosystems) with the following protocol: 50°C for 2 min, 95°C for 2 min, 40 cycles of 95°C for 15 sec and 65°C for 60 sec, with a final melt curve (95°C for 15 sec, 60°C for 60 sec). Ct values were calculated using the ThermoFisher qPCR Design and Analysis Platform. Relative expression was determined by calculating the ΔCt of the target gene normalized to a housekeeping gene. All samples were run in technical triplicate.

#### 16S Differential Abundance

Relative abundances were calculated for each community composition from 16S counts data, normalizing individual species counts to total counts in a sample. Principal component analysis of differential abundances on Days 0 and 7 was done in Prism, with the following parameters: Standardize (scale data to have mean of 0 and SD of 1) and Selecting PCs with eigenvalues greater than 1 (Kaiser rule). At Day 7, taxa were first filtered by Kruskal-Wallis test (α = 0.05) and Mann-Whitney U test (α = 0.05) between DSS and Healthy groups. Effect size was quantified using a univariate LDA score (log_10_(1 + |Mean_DSS_ − Mean_Healthy_|) computed on relative abundances scaled to 1 × 10⁶, consistent with the internal scaling convention of LEfSe. Differentials were considered significant if they passed both statistical filters and had a |Log_2_Fold Change| ≥ 1.0.

#### RNA-seq Differential Expression

Sequencing reads were quality filtered (FastQC^96^) and trimmed (Trimmomatic^97^), they were aligned to the DMC reference genomes (Bowtie2^98^). Aligned reads counts were binned to reference annotations using featureCounts (Rsubread^99^) and tested for differential expression (DESeq2^100^) species by species to control for abundance differences. Low read count, rRNA, and tRNA genes were filtered out of counts files before DESeq2. Differential expression analysis directly compared conditions, generating lists of significantly up- and down-regulated genes (P_adj_ < 0.05). *In vitro* vs *in vivo* analysis used a threshold of |Log_2_Fold Change| > 2 and DSS vs Healthy used a threshold of |Log_2_Fold Change| > 1 to include more candidates for CSS identification. DMC community genomes were further annotated via PATRIC^101^ to add SEED Subsystems and SEED subclasses, KEGG^102^ to add KEGG pathways, and PROKKA^103^ to add PFAM annotations. Enrichment analysis was performed using the GSEA() function from the clusterProfiler package^104^ to search for pathways and subsystems significantly enriched (P_adj_ < 0.05) between the two cell types. All analysis was done in Rstudio and plots generated using the tidyverse R package^105^. Claude (Anthropic) was used to assist in writing R code for data analysis and plot generation; all code was reviewed and validated by the authors.

#### BCSeq Analysis

UniqueSeq analysis from Genewiz delivered a counts matrix of binned unique sequences. A custom R script was used to filter these sequences by size, extract all 9 bp barcodes that matched to registered CSS and located immediately upstream of the expected NanoLuc sequence, and quantify by barcode. Raw barcode counts for DNA and RNA libraries were tabulated for each construct (CSS) and its associated barcodes. DNA count distributions for each species were plotted to confirm even library representation. RNA and DNA counts for each sample were normalized to total library size (counts divided by column sum), and barcode-level relative expression (RelExp) was calculated as the ratio of normalized RNA to normalized DNA counts for each barcode in each sample. Low count RNA and DNA Barcodes were filtered out for each Pool, depending on sequencing read depth and the number of CSS in the Pool. Correlation between barcodes for the same CSS was assessed by Pearson correlation of log2 transformed RelExp values, across all samples combined and for each sample individual. CSS RelExp values were averaged across the two barcodes, median-normalized, and log2-transformed. Fold change between conditions was calculated as the ratio of mean RelExp across condition 1 (i.e. DSS) to mean RelExp across condition 2 (i.e. Healthy), and constructs with |log_2_FC| ≥ 1 were flagged as differential. For heatmap plotting, rows were ordered by fold change and columns hierarchically clustered within each condition. CSS activity for each species library was also assessed using MPRAnalyze^64^ to perform library-size normalization and formal statistical tests for differential activity. CSSs with more than 2-fold differential expression and P_adj_ < 0.05 (Benjamini-Hochberg) were classified as upregulated. Final Log_2_FC values were compared to RNA-seq DESeq2 differential expression using a Pearson correlation calculation. Bt and Pv CSS libraries were run independently for all analysis, then combined for plotting purposes. All analysis and plots were done in Rstudio, using the tidyverse^105^ and MPRAnalyze^64^ package. Claude (Anthropic) was used to assist in writing R code for data analysis and plot generation; all code was reviewed and validated by the authors.

#### CSS Selection (SSMiner)

For Pools 1 and 2 a computational decision tree (SSMiner) was designed to select CSS from differential expression data based off of the following criteria: (1) the gene was upregulated, (2) the gene is part of a multigene operon (Rockhopper mapping), and either (3) the gene is located near a regulator or (4) is a regulator. Differentially expressed genes (DEGs) within five genes of a designated regulator in either direction were selected as CSS candidates, along with any differentially expressed regulators themselves (DEGREGS). Regulators were defined as genes annotated as transcription factors, sigma factors, two-component system components, response regulators, activators, or DNA-binding proteins, as well as selected PFAM clans CL0123, CL0089, CL0108, CL0144, CL0177, CL0304, CL0319, CL0439, CL0548, and CL0654. For Pool3, SSMiner was adjusted to filter DEGS for baseline expression (baseMean ≥ 10) and classify GSEA involvement (KEGG, PATRIC Subclass, and PATRIC Subsystem). Regulatory context was determined using product-name keyword matching (e.g., LuxR, LysR, AraC, GntR, TetR, HTH, sigma factor, two-component system) and a more restricted regulator Pfam domain list (PF00196, PF00486, PF13185). CSS were then ranked according to differential expression, GSEA involvement, operon membership and regulatory context. DEGs sharing a first gene were consolidated into a single promoter candidate and DEGREGS separated into their own list. Multiple downregulated CSS were selected as well to balance the dynamic range of the Pool. Lists of selected Pool1, Pool2, and Pool3 CSS are in Table S11. Claude (Anthropic) was used to assist in writing R code; all code was reviewed and validated by the authors.

#### Data Analysis and Statistics

Statistical analyses were performed using GraphPad Prism version 10.5.0, Microsoft Excel, and R/Rstudio. The test details are included in the main text and figure legends.

#### Key Resources Table

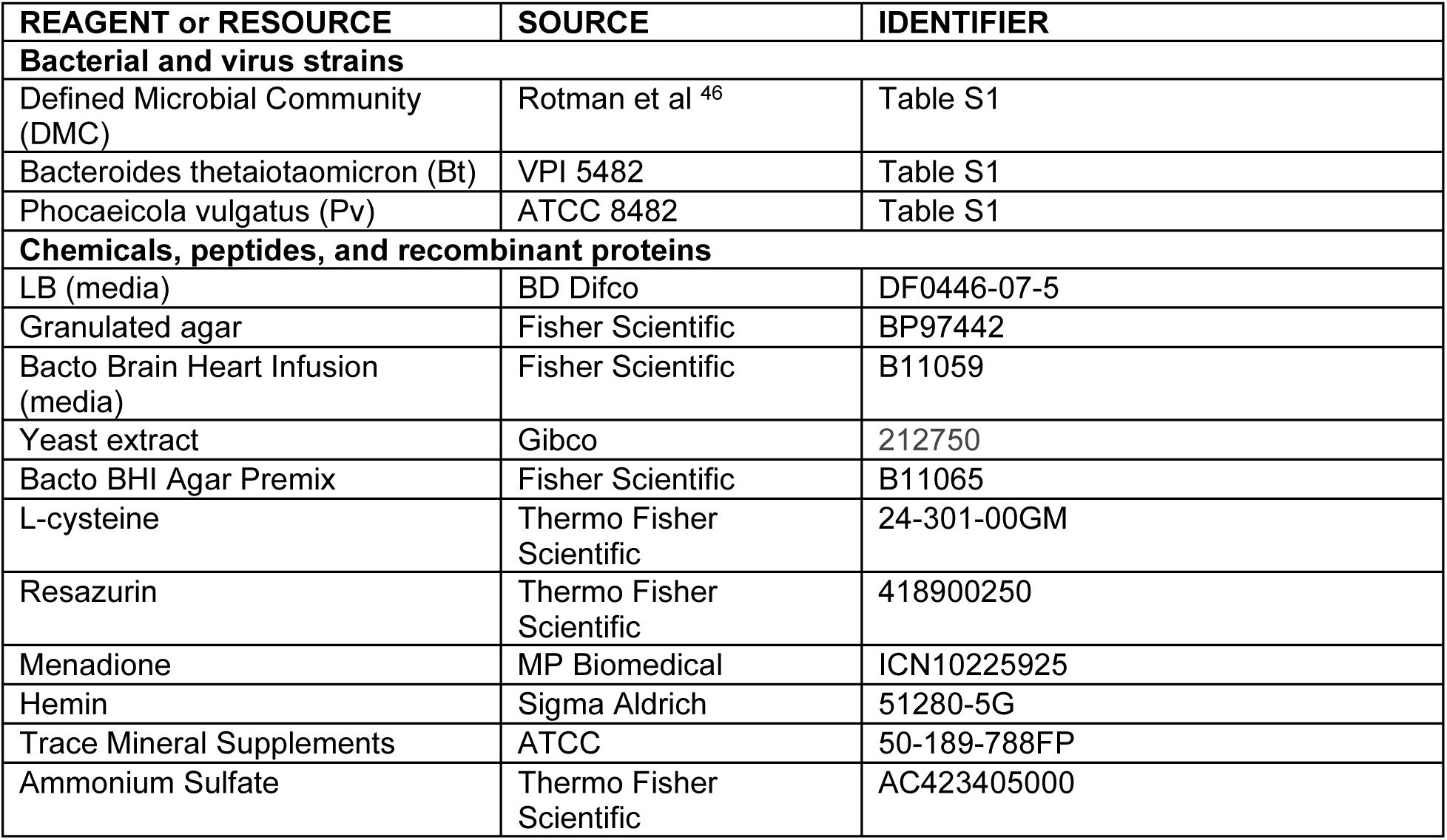
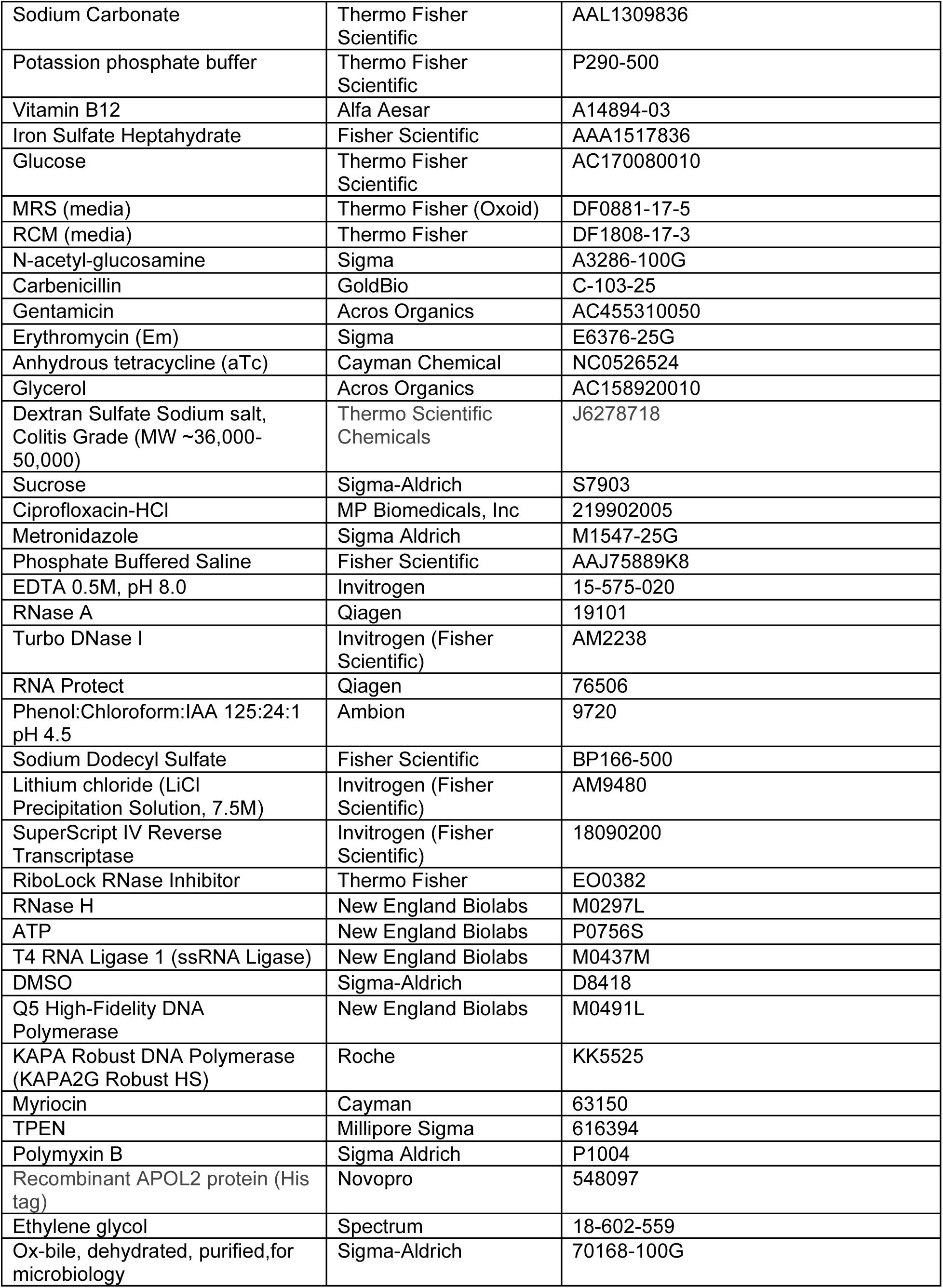
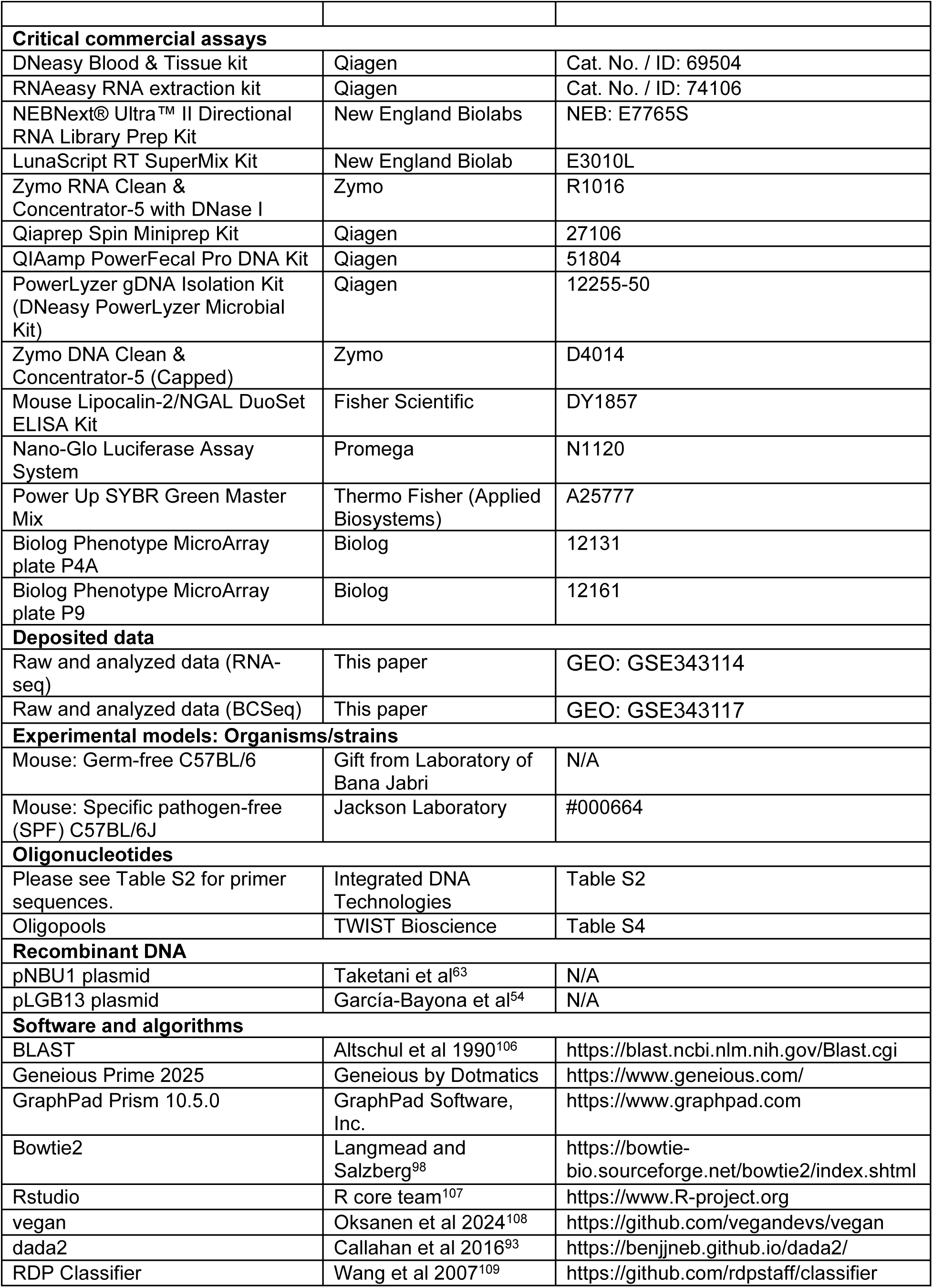
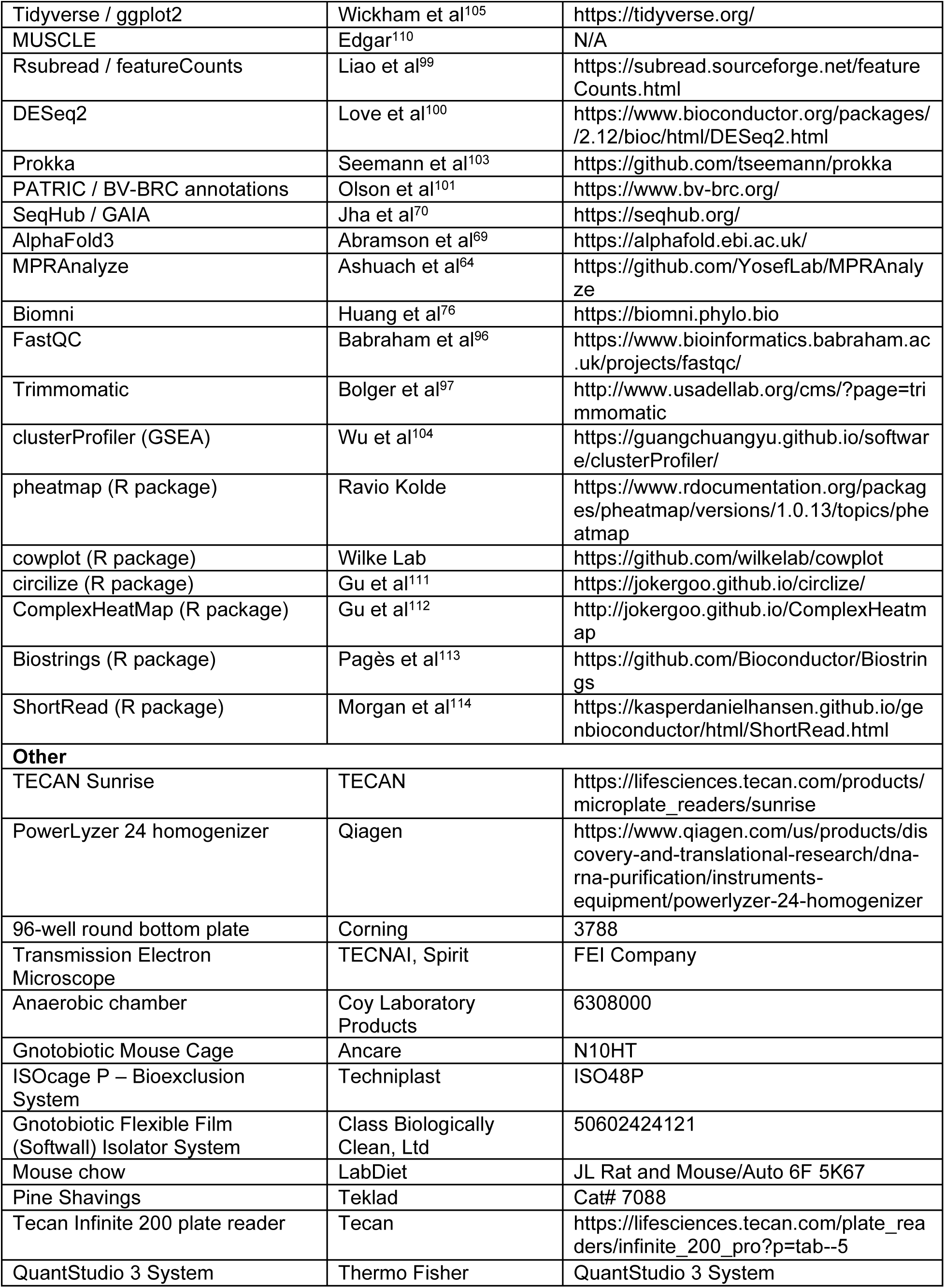
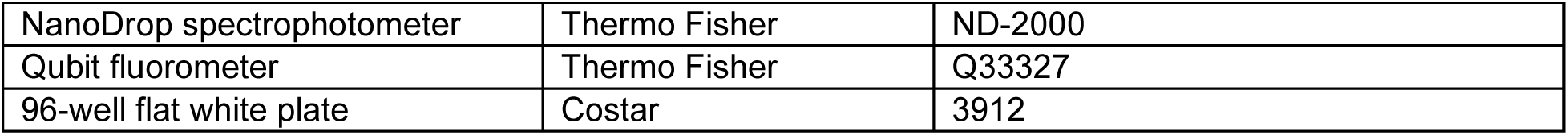

## TABLES

1. *Table S1, List of Strains Used in This Study, related to STAR Methods.*
2. *Table S2, List of Primers Used in This Study, related to STAR Methods*
3. *Table S3, List of Plasmids Used in This Study*
4. *Table S4, List of Promoter Sequences Used in This Study, related to STAR Methods*
5. *Table S5, List of Deletion Mutant Segments Used in This Study*
6. *Table S6, Defined Microbial Community Expanded Annotations*
7. *Table S7, In vivo vs in vitro (VV) transcriptomics results (RNA-seq, DESeq2)*
8. *Table S8, Gene Set Enrichment Analysis (GSEA) of VV RNA-seq*
9. *Table S9, DSS vs Healthy (DSS) transcriptomics results (RNA-seq, DESeq2)*
10. *Table S10, Gene Set Enrichment Analysis (GSEA) of DSS RNA-seq*
11. *Table S11, Candidate Sensor Systems (CSS) selections, annotations, and Pool involvement*
12. *Table S12, Pool1 DSS BCSeq Results*
13. *Table S13, Pool2 DSS BCSeq Results*
14. *Table S14, Pool2 VV BCSeq Results*
15. *Table S15, Pool3 DSS BCSeq Results*
16. *Table S16, Top Candidate Sensor Systems (CSSs) selected for in vitro analysis*
17. *Table S17, BT32 KO Mutants RNA-seq Results (ΔBT_4647 vs WT)*
18. *Table S18, BT32 KO Mutants RNA-seq Results (ΔBT_4648-49 vs WT)*

