## Supplemental Data for "Mining Microbial Transcriptomes to Engineer Cell-Based Bacterial Biosensors in Gut-Resident Bacteroidaceae"

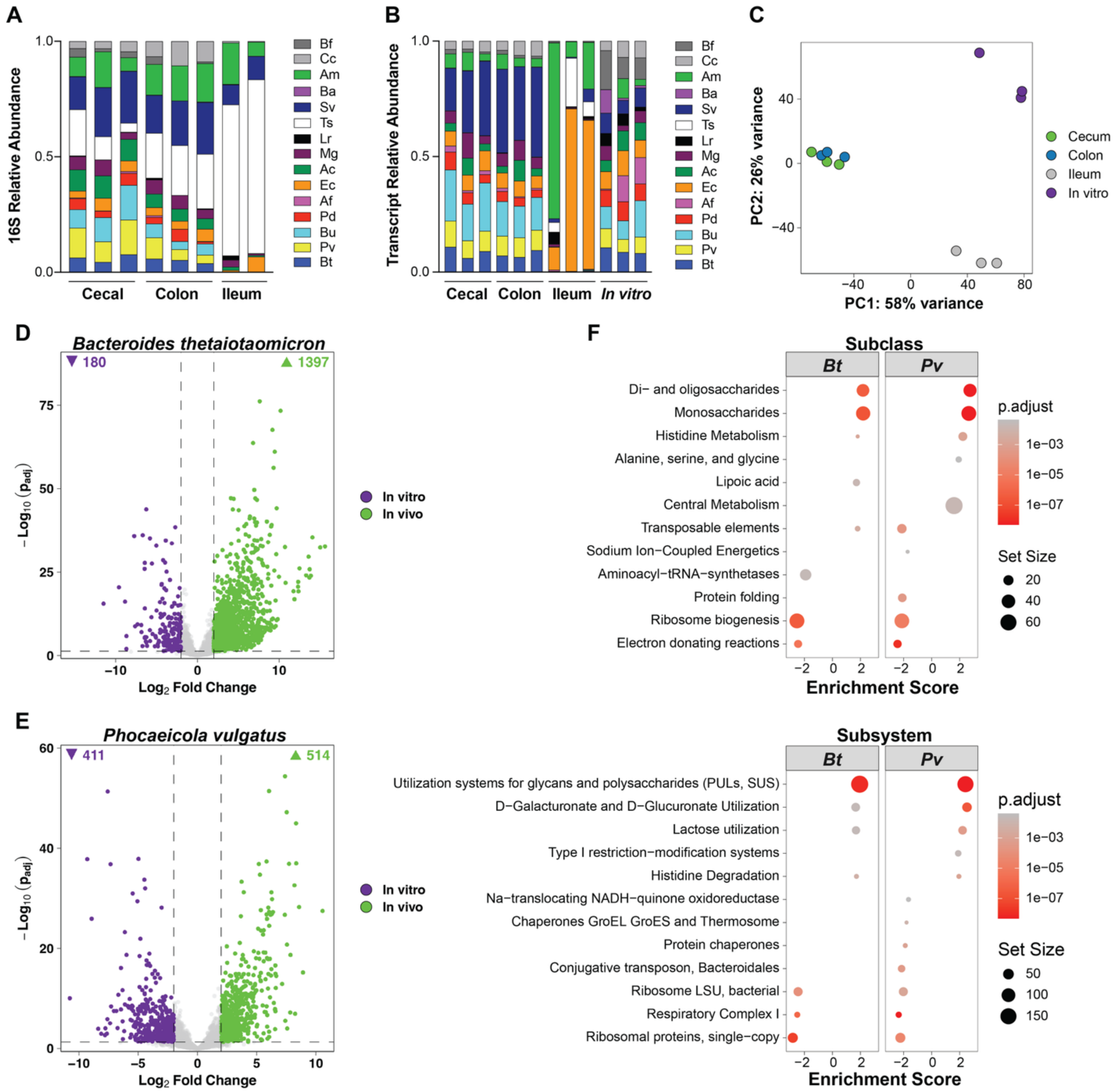


<u>Supplemental Figure 1:</u> Community composition and transcriptional profiling of Defined Microbial Community (DMC) in *in vivo* and *in vitro*, with differential expression and pathway analysis of *Bacteroides thetaiotaomicron* and *Phocaeicola vulgatus*

(A) Relative abundance of DMC from 16S rRNA sequencing of *in vivo* cecal, colon, and ileum samples (n = 3 per condition, Ileum n = 2)

(B) Transcript relative abundance of DMC from RNA-sequencing of *in vivo* cecal, colon, and ileum contents, and *in vitro* mid-log growth cultures (n = 3 per condition)

(C) Principal component analysis of RNA-seq data from cecum, colon, ileum, and *in vitro* conditions.

(D) Volcano plot of differential expression analysis (DESeq2) of *Bacteroides thetaiotaomicron* ($Log_2$Fold Change > 2, $P_{adj} < 0.05$) comparing *in vitro* to *in vivo* (cecal) conditions

(E) Volcano plot of differential expression analysis of *Phocaiecola vulgatus* ($Log_2$Fold Change > 2, $P_{adj} < 0.05$) comparing *in vitro* to *in vivo* (cecal) conditions

(F) Gene Set Enrichment Analysis (GSEA) of *Bacteroides thetaiotaomicron* (Bt) and *Phocaeicola vulgatus* (Pv) comparing *in vitro* to *in vivo* (cecal) conditions. GSEA analysis on SEED Subclass and Subsystems, from PATRIC BV-BRC annotations.

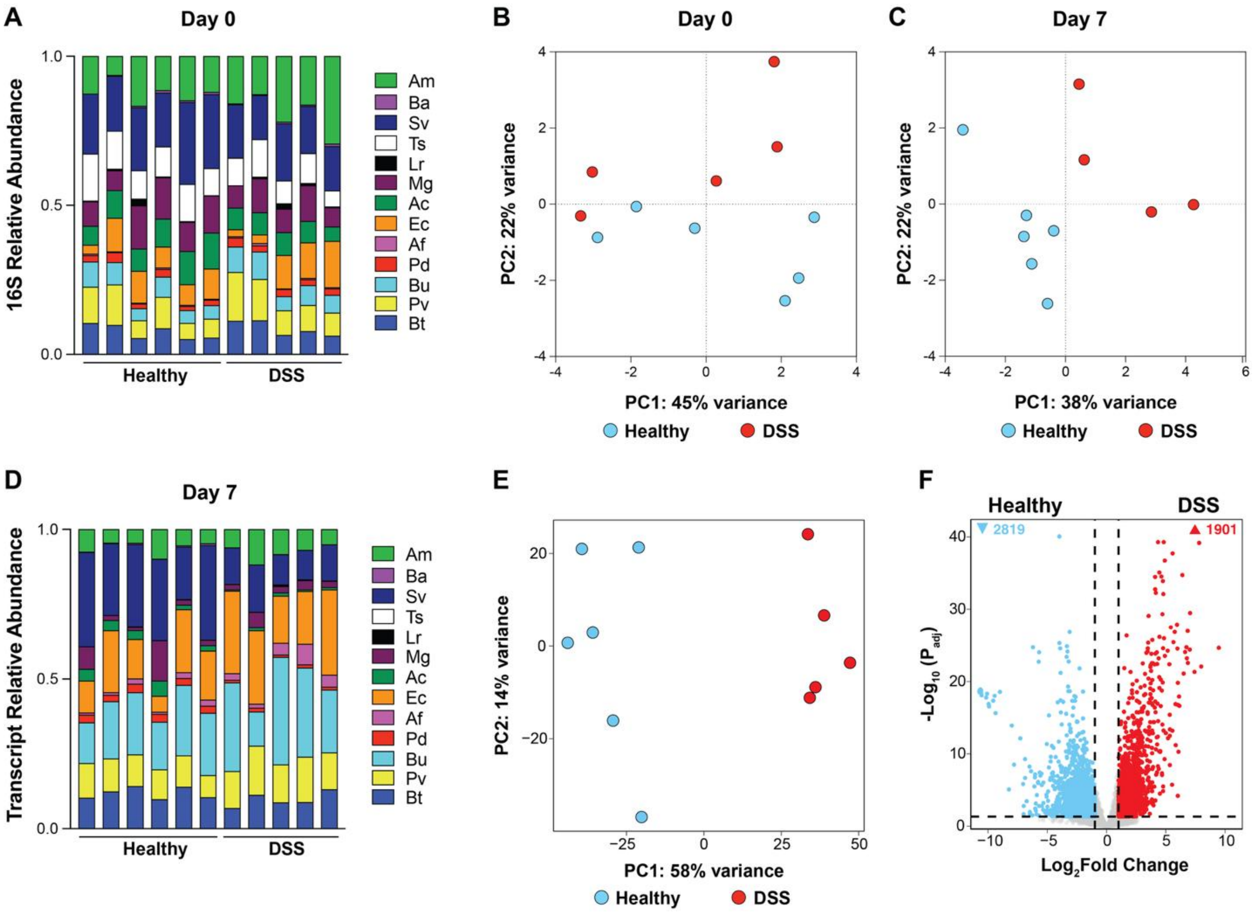


Supplemental Figure 2: Community composition and transcriptional profiling of Defined Microbial Community in DSS versus Healthy conditions

(A) Relative abundance of DMC from 16S rRNA sequencing of DSS and Healthy conditions on Day 0, before administration of 2.5% DSS drinking water (n = 6 per condition, 1 DSS samples not sampled)
(B) Principal component analysis (Prism) of Day 0 16S rRNA sequencing Data. No clear separation between conditions at the start of the DSS treatment
(C) Principal component analysis (Prism) of Day 7 16S rRNA sequencing Data. Separation between conditions at the end of the DSS treatment
(D) Transcript relative abundance of DMC from RNA-sequencing of Healthy and DSS cecal contents (n = 6 per condition, one DSS ID dropped from sequencing)
(E) Principal component analysis of variance-stabilizing transformed counts from DESeq2 analysis. Clear separation between Healthy and DSS groups.
(F) Volcano plot of differential expression analysis (DESeq2) of all genes in DMC ($Log_2$Fold Change > 1, $P_{adj} < 0.05$) comparing DSS to Healthy conditions.

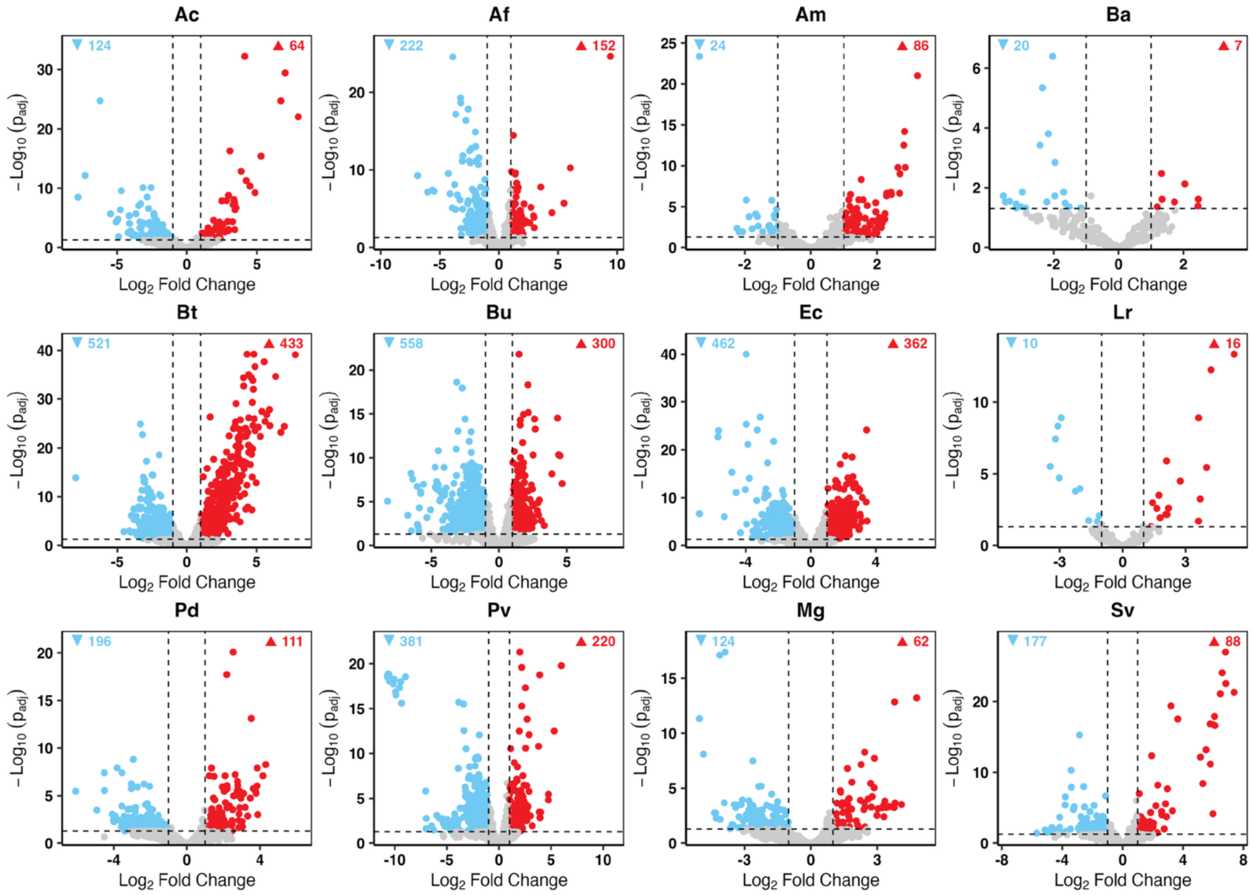


Supplemental Figure 3: Differential expression of individual DMC members in DSS versus Healthy conditions. Individual volcano plots of differential expression analysis (DESeq2) of each individual DMC member ($|Log_2Fold\ Change| > 1$, $P_{adj} < 0.05$). Full RNA-seq differential expression analysis in Table S9.

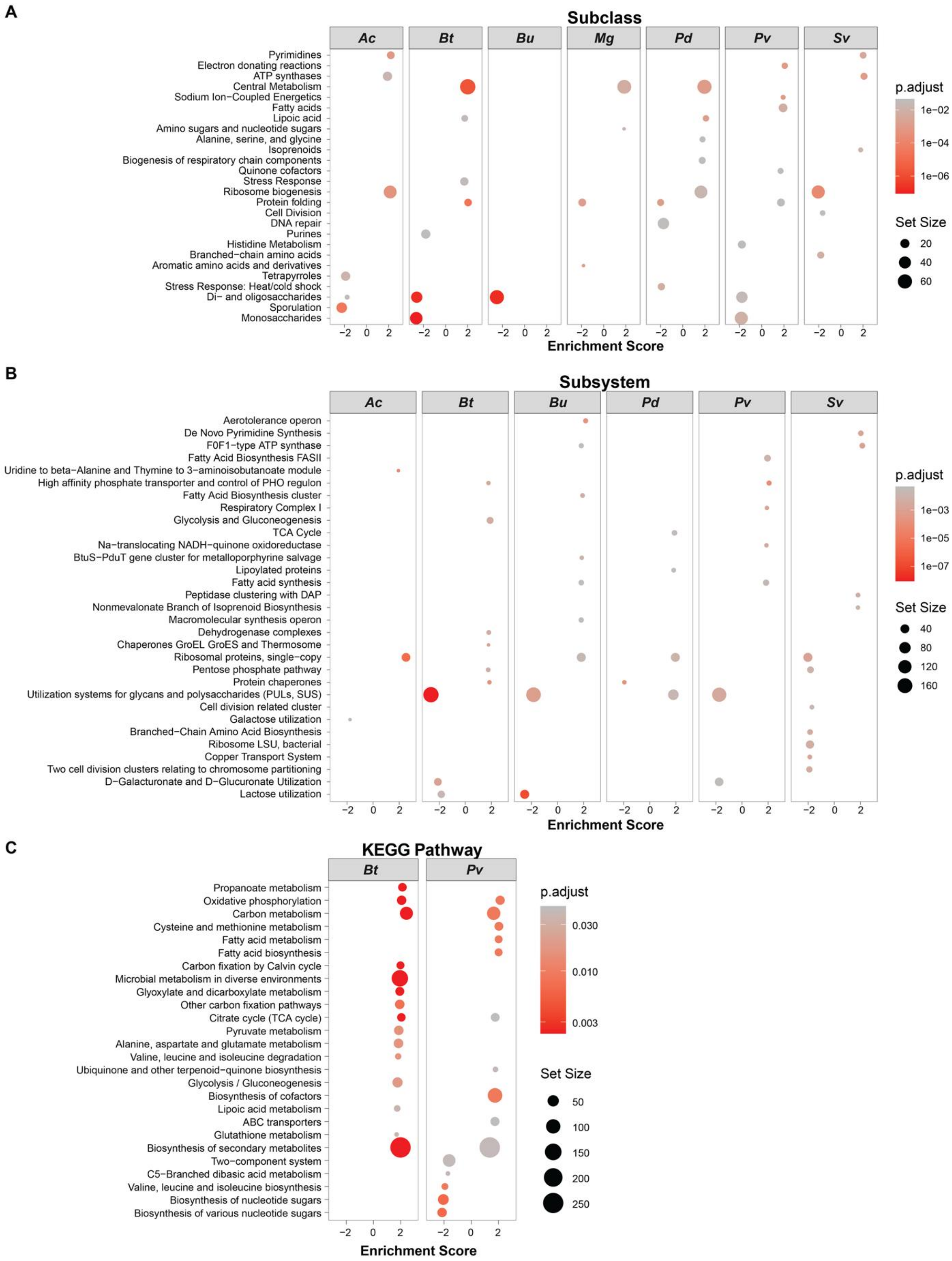


<u>Supplemental Figure 4:</u> Gene Set Enrichment Analysis (GSEA) of DSS vs Healthy RNA-seq shows a community under general stress and metabolic change

(A) Gene set enrichment analysis (GSEA) using clusterProfiler for PATRIC annotated SEED Subclass from DSS vs Healthy results (DESeq2) in *Anaerostipes caccae (Ac), Bacteroides thetaiotaomicron (Bt), Bacteroides uniformis (Bu), Mediterraneibacter gnavus (Mg), Parabacteroides distasonis (Pd), Phocaiecola vulgatus (Pv), Subdoligranulum variabile (Sv)*. Set Size > 5, $P_{adj} < 0.05$.
(B) GSEA of SEED Subsystems from DSS vs Healthy results (DESeq2) in Ac, Bt, Bu, Pd, Pv, and Sv. Set Size > 5, $P_{adj} < 0.05$.
(C) GSEA of KEGG pathways from DSS vs Healthy results (DESeq2) in Bt and Pv. Set Size > 5, $P_{adj} < 0.05$.

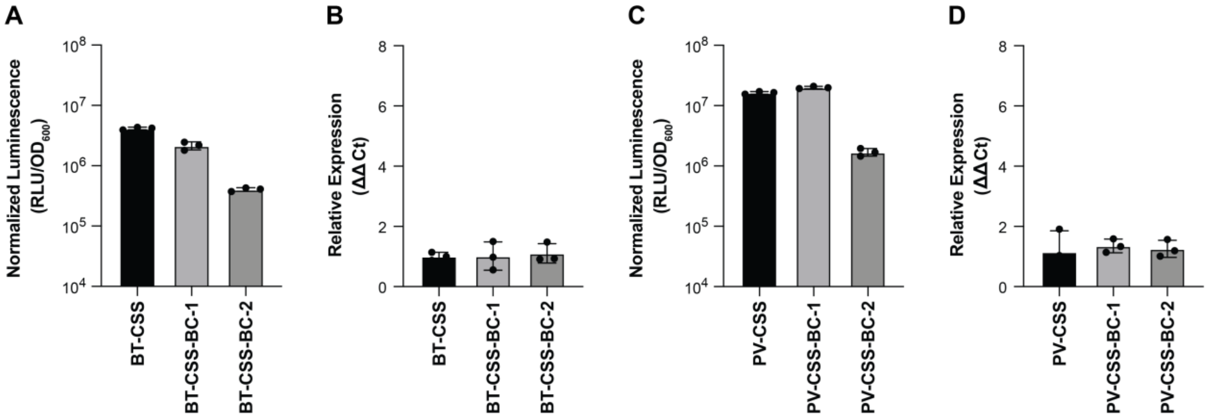


Supplemental Figure 5: Barcoded Nanoluciferase (NL) reduces CSS expression via luminescence but does not inhibit transcription of NL RNA in both Bt and Pv CSS.

(A) Normalized Luminescence from overnight cultures of a representative BT-CSS and two barcoded BT-CSS expression vectors. BT-CSS-BC-2 dropped 10.3X in normalized luminescence compared to no barcode control BT-CSS.
(B) Relative Expression (ΔΔCt) of BT-CSS and barcoded variants measured by qPCR. ΔΔCt calculated using Ct values from NL and Bt *gmk* housekeeping gene.
(C) Normalized Luminescence from overnight cultures of a representative PV-CSS and two barcoded BT-CSS expression vectors. BT-CSS-BC-2 dropped 9.6X in normalized luminescence compared to no barcode control BT-CSS.
(D) Relative Expression (ΔΔCt) of PV-CSS and barcoded variants measured by qPCR. ΔΔCt calculated using Ct values from NL and Pv *gyrB* housekeeping gene.

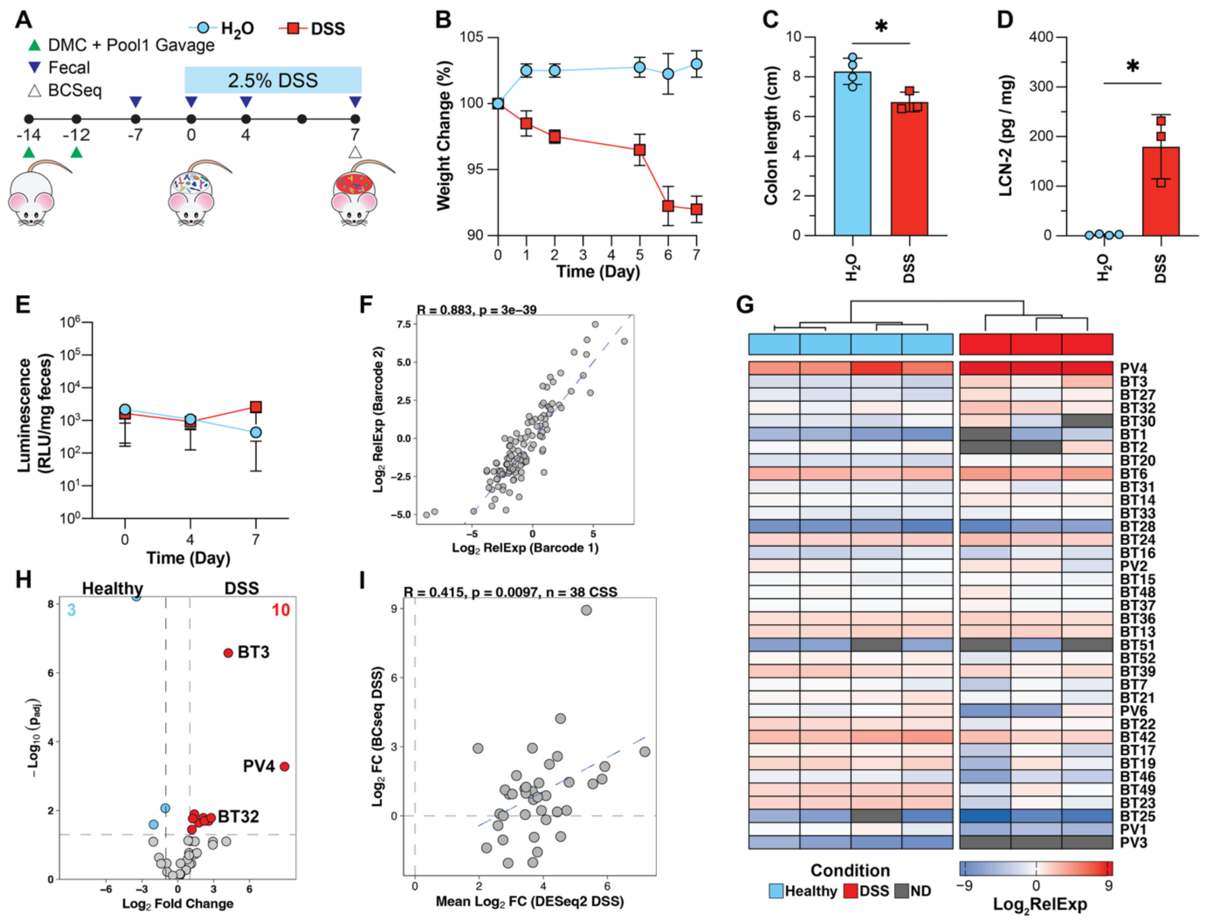


Supplemental Figure 6: *In vivo* screening of CSS Pool1 in a gnotobiotic DSS colitis model identifies responsive CSS in Bt and Pv

(A) Schematic of CSS Pool DSS Timeline. DMC + CSS Pool1 are gavaged together, twice, at least two days apart and at least two weeks before DSS administration. Engraftment of CSS and DMC members is checked one week before DSS administration. Fecals are collected on Days 0 and 4, and Cecal contents collected for Barcode Sequencing (BCSeq) analysis on Day 7.

(B) DSS group loses weight over time compared to $H_2O$ control groups, as a measure of percent change in body weight relative to baseline (n = 4 $H_2O$, 3 DSS)

(C) Colon length (cm) in $H_2O$ versus DSS groups

(D) Lipocalin-2 (LCN-2) concentrations (pg/mg of feces) in $H_2O$ versus DSS groups

(E) CSS Pool luminescence (RLU / mg of feces) in $H_2O$ vs DSS groups.

(F) Correlation of $log_2$RelExp of Pool1 CSS repeat barcodes. Pearson correlation.

(G) Heatmap of Pool1 RelExp per CSS per mouse ID. Rows are sorted top to bottom by RelExp Fold change (DSS to Healthy) and hierarchal clustering determined column order. Barcodes not detected (ND) are colored gray.

(H) MPRAnalyze results of CSS Pool1 fold change in DSS versus Healthy. 10 CSS were significantly upregulated ($Log_2$Fold Change > 1, $P_{adj} < 0.05$, Benjamini-Hochberg) and 3 CSS significantly downregulated ($Log_2$Fold Change < -1, $P_{adj} < 0.05$, Benjamini-Hochberg)

(I) Comparison of BCSeq (Log2FC) and RNA-seq results (Log2FC) for CSS Pool1. Pearson correlation.

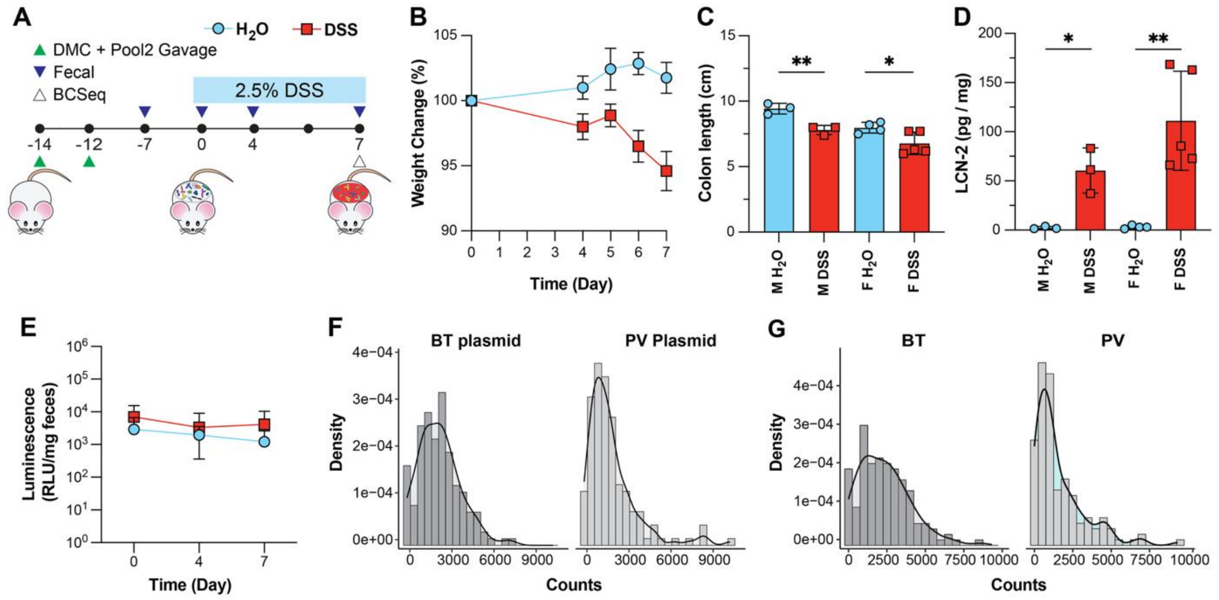


Supplemental Figure 7: DSS colitis model results and CSS Pool2 library distribution from *in vivo* screening

(A) Schematic of CSS Pool2 DSS Timeline
(B) DSS group loses weight over time compared to $H_2O$ control groups, as a measure of percent change in body weight relative to baseline (n = 7 $H_2O$, 8 DSS)
(C) Colon length (cm) in $H_2O$ versus DSS groups in Male and Female mice
(D) Lipocalin-2 (LCN-2) concentrations (pg/mg of feces) in $H_2O$ versus DSS groups in Male and Female mice
(E) CSS Pool luminescence (RLU / mg of feces) in $H_2O$ vs DSS groups.
(F) Histogram of unique barcode counts assigned to Pool2 CSSs for *E. coli* S17 plasmid libraries for Bt and Pv. Only exactly barcode matches are plotted.
(G) Histogram of unique barcode counts assigned to Pool2 CSSs for conjugated Bt and Pv gDNA libraries. Only exactly barcode matches are plotted.

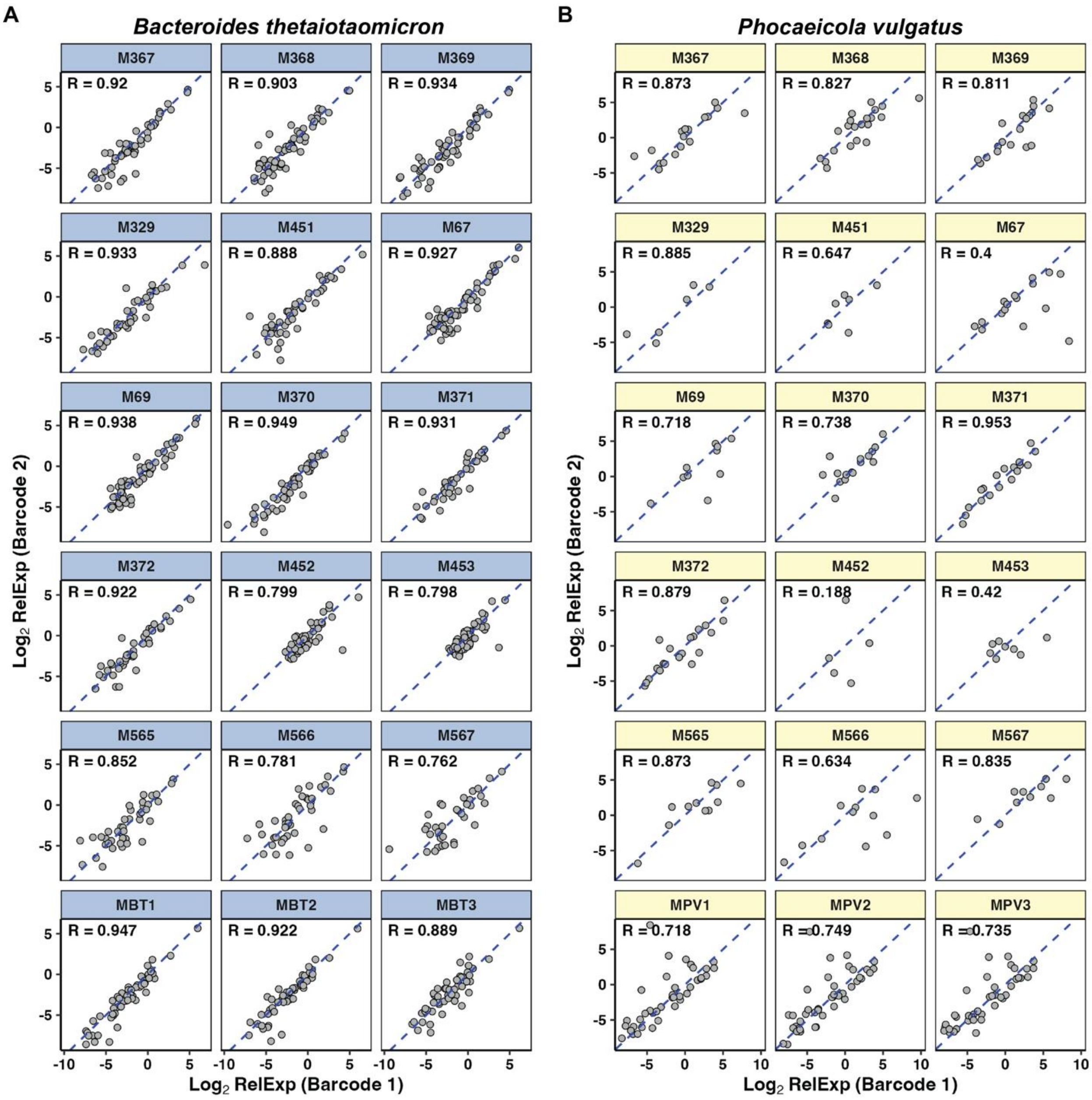


Supplementary Figure 8: Repeated barcodes per CSS in Pool2 show high degree of correlation in relative expression across individual healthy, DSS, and *in vitro* samples

(A) Correlation of $\log_2$RelExp of Barcode 1 vs Barcode 2, for each Pool2 Bt CSS in individual samples across Healthy, DSS and *in* vitro conditions. Pearson correlation.

(B) Correlation of $\log_2$RelExp of Barcode 1 vs Barcode 2, for each Pool2 Pv CSS in individual samples across Healthy, DSS and *in* vitro conditions. Pearson correlation.

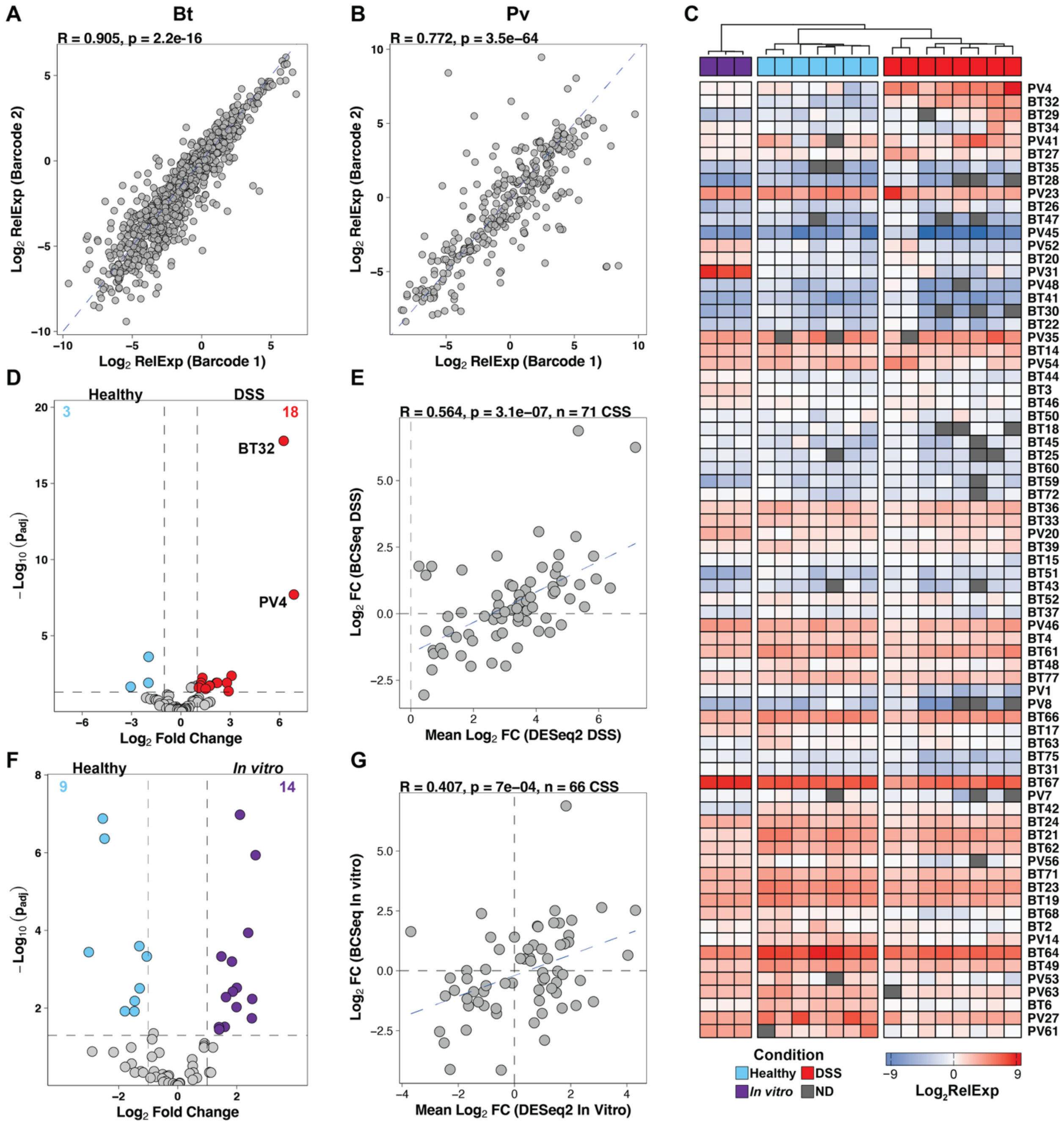


Supplementary Figure 9: *In vivo* and *in vitro* screening of CSS Pool2 in healthy, disease, and mid-log growth conditions passes key correlation and QC metrics and identifies responsive CSS in Bt and Pv

(A) Correlation of $log_2$RelExp of Barcode 1 vs Barcode 2, for all Pool2 Bt CSS in all samples across Healthy, DSS and *in* vitro conditions. Pearson correlation.

(B) Correlation of $log_2$RelExp of Barcode 1 vs Barcode 2, for all Pool2 Pv CSS in all samples across Healthy, DSS and *in* vitro conditions. Pearson correlation.

(C) Heatmap of Pool2 RelExp per CSS per mouse ID across Healthy, DSS and *in vitro* conditions. Rows are sorted top to bottom by RelExp Fold change (DSS to Healthy) and hierarchal clustering determined column order. Barcodes not detected (ND) are colored gray.

(D) MPRAnalyze results of CSS Pool2 fold change in DSS versus Healthy. 18 CSS were significantly upregulated ($Log_2$Fold Change > 1, $P_{adj} < 0.05$, Benjamini-Hochberg) and 3 CSS significantly downregulated ($Log_2$Fold Change < -1, $P_{adj} < 0.05$, Benjamini-Hochberg). BT32 and PV4 were the top CSS in DSS conditions.

(E) Comparison of BCSeq (Log2FC) and RNA-seq results (Log2FC) for CSS Pool2. Pearson correlation, 71 CSS detected and measured.

(F) MPRAnalyze results of CSS Pool2 fold change in *In vitro* versus Healthy. 14 CSS were significantly upregulated *in vitro* ($Log_2$Fold Change > 1, $P_{adj} < 0.05$, Benjamini-Hochberg) and 9 CSS were significantly upregulated in Healthy mice gut microbiomes ($Log_2$Fold Change < -1, $P_{adj} < 0.05$, Benjamini-Hochberg).

(G) Comparison of BCSeq (Log2FC) and RNA-seq results (Log2FC) for *in vitro* CSS Pool2. Pearson correlation, 66 CSS detected and measured.

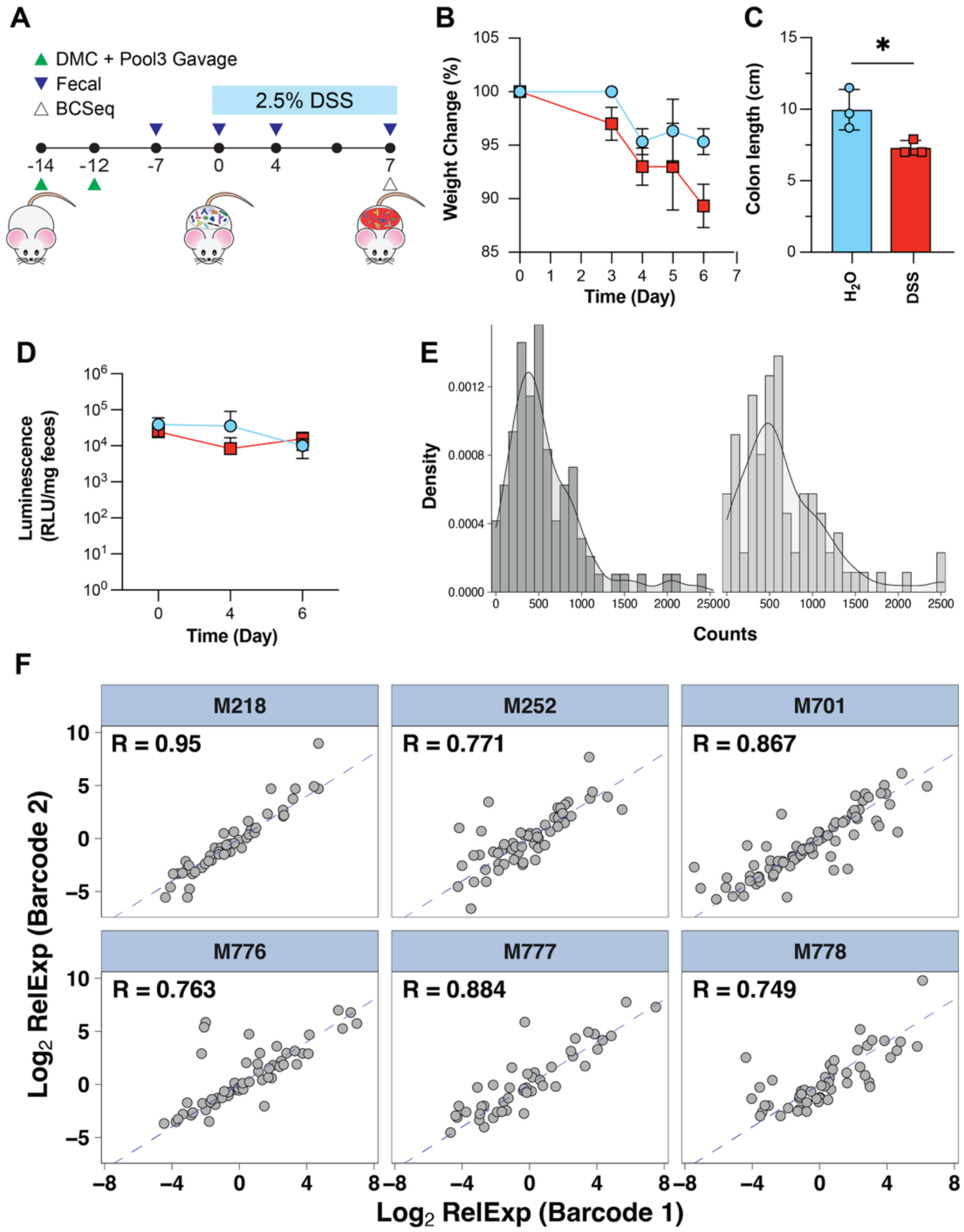


<u>Supplementary Figure 10:</u> DSS colitis model results, CSS Pool3 library distribution, and individual sample repeated barcode correlation from *in vivo* screening

(A) Schematic of CSS Pool3 DSS Timeline
(B) DSS group loses weight over time compared to $H_2O$ control groups, as a measure of percent change in body weight relative to baseline (n = 3 $H_2O$, 3 DSS)
(C) Colon length (cm) in $H_2O$ versus DSS groups in Male and Female mice
(D) CSS Pool luminescence (RLU / mg of feces) in $H_2O$ vs DSS groups.
(E) Histogram of unique barcode counts assigned to Pool3 CSSs for conjugated Bt and Pv gDNA libraries. Only exactly barcode matches are plotted.
(F) Correlation of $log_2$RelExp of Barcode 1 vs Barcode 2, for all Pool2 CSS (Bt and Pv) in individual samples across Healthy and DSS conditions. Pearson correlation.

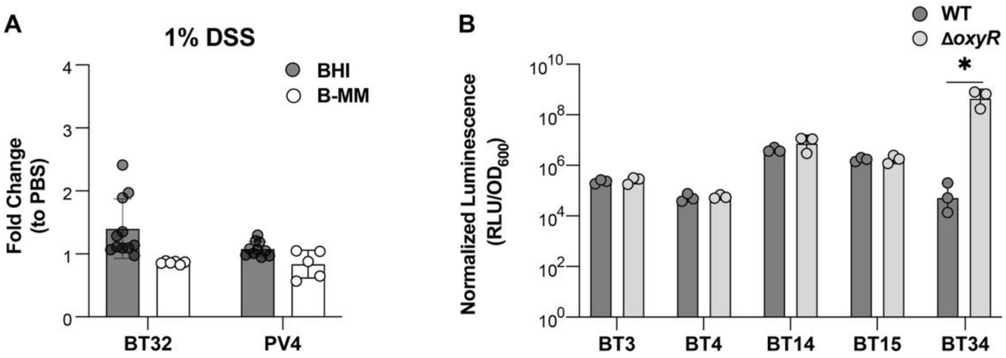


Supplementary Figure 11: *Top CSS in vitro response to DSS in multiple medias and deletion of Bt oxyR gene*

(A) BT32 and PV4 have minimal response to 1% DSS in BHI and BMM. BT32 and PV4 subcultures were grown to mid-log phase and stimulated with 1% DSS or PBS, then measured two hours post induction. Fold Change calculated by dividing DSS normalized luminescence by PBS normalized luminescence

(B) Expression (Normalized Luminescence) of top CSS in wild-type (WT) and *oxyR* deletion mutant strains. Overnight cultures in BHIS media

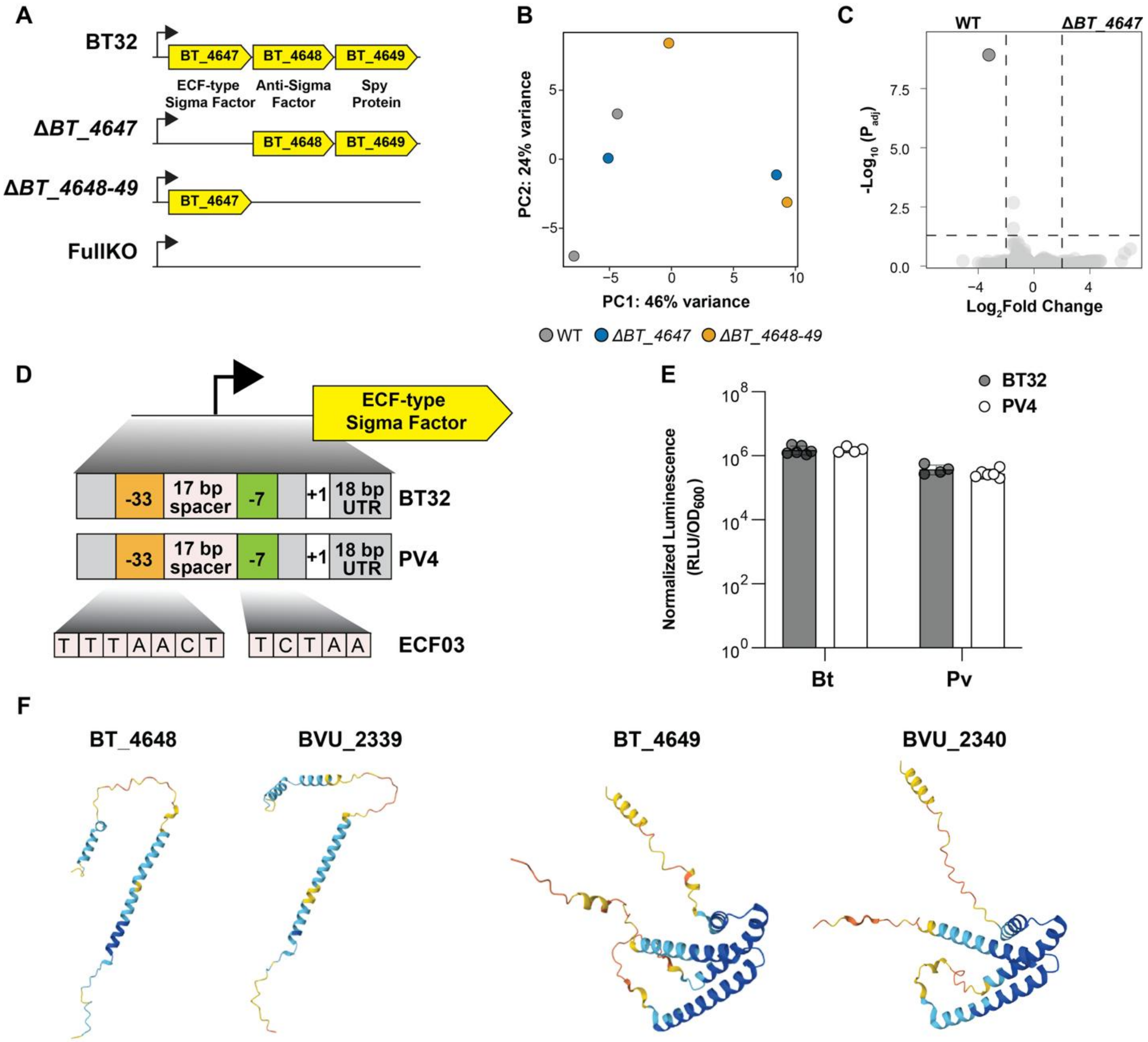


Supplementary Figure 12: Characterization of homologous BT32 and PV4 operons shows tight regulons, identical promoter architecture, and functionality across chassis

(A) Schematic of BT32 deletion mutant strains
(B) Principal component analysis (PCA) of variance stabilizing transform RNA seq results for wild-type (WT), deletion of sigma factor (BT_4647), and deletion of two downstream genes (BT_4848, BT4849). No separation between groups from PCA analysis (Prism)
(C) Volcano plot of differential expression analysis (DESeq2) comparing wild-type Bt (WT) to the Bt mutant strain with the sigma factor deleted (BT_4647). The WT upregulation gene is BT_4647 (gray).
(D) Promoter architecture of BT32 and PV4. Promoter has an 18 base pair 5' untranslated region (UTR), with identical -7/-33 sites and spacer size. ECF-type sigma factor is annotated as a ECF03 (Rhodius paper I think).
(E) Expression of BT32 and PV4 in multiple chassis. Normalized expression shows CSS expression is host chassis dependent. (n = 6)

(F) Alphafold3 predicted structures for BT_4648 / BVU_2339 and BT_4649 / BVU_2340. Structures are highly similar.

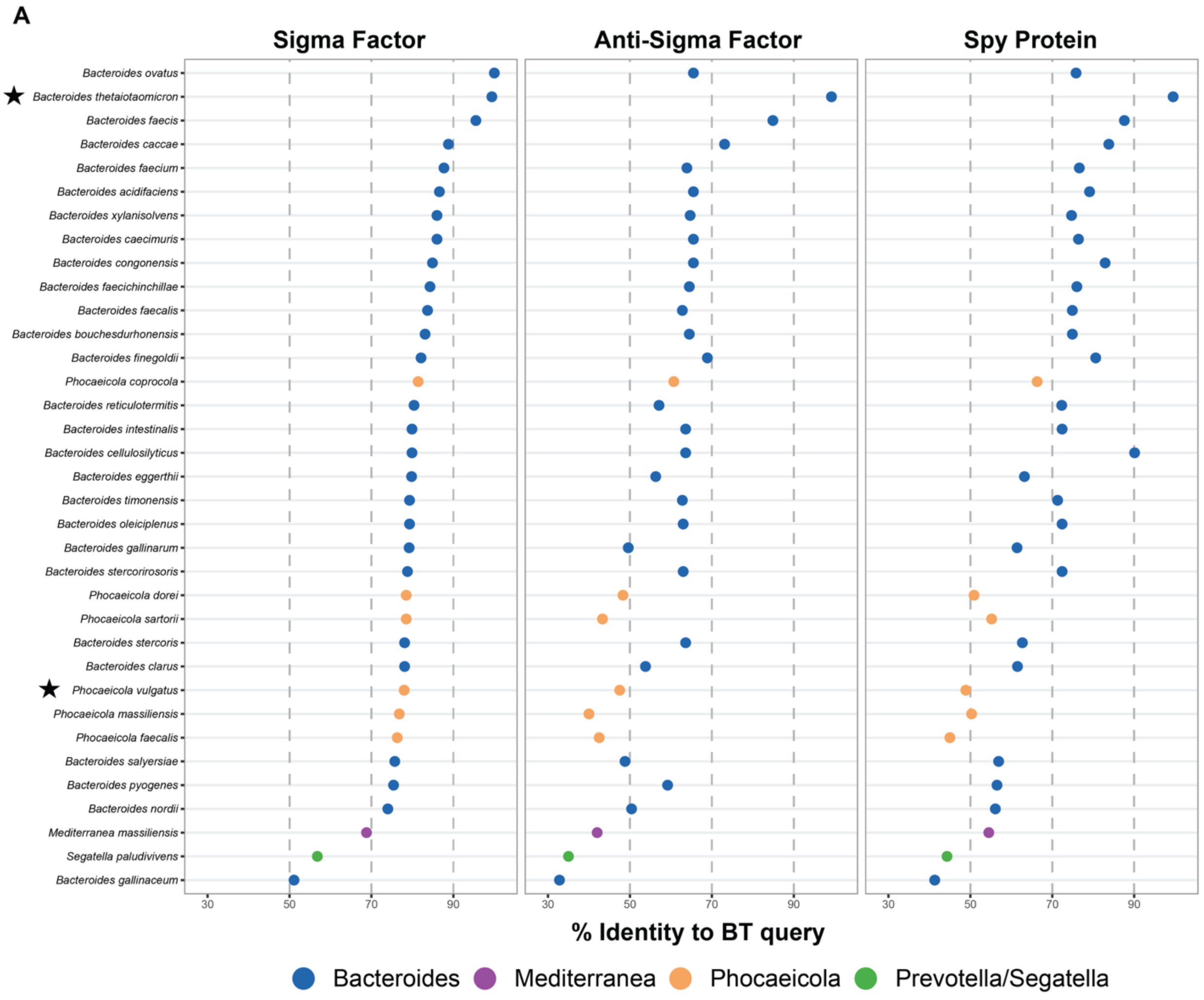


Supplementary Figure 13: BT32 phylogeny and synteny analysis show broad conservation across Bacteroidales

(A) Gene synteny analysis of BT32 three gene operon. Each protein (Sigma Factor, Anti-Sigma Factor, Spy Protein) was used as a query in BLAST sequence similarity searches, then ordered according to percent identify to original query, and syntenic operon organization. Of the 35 organisms with detectable similarity to all three genes, 34 carry all three genes together in the same genomic location and in the same order. Point color indicates which genus or family the organism belongs to. The operon is found predominantly in gut-associated Bacteroides and Phocaeicola species, with more divergent copies in distantly related Bacteroidales lineages including Prevotella, Segatella and Mediterranea. Stars indicate Bt and Pv.

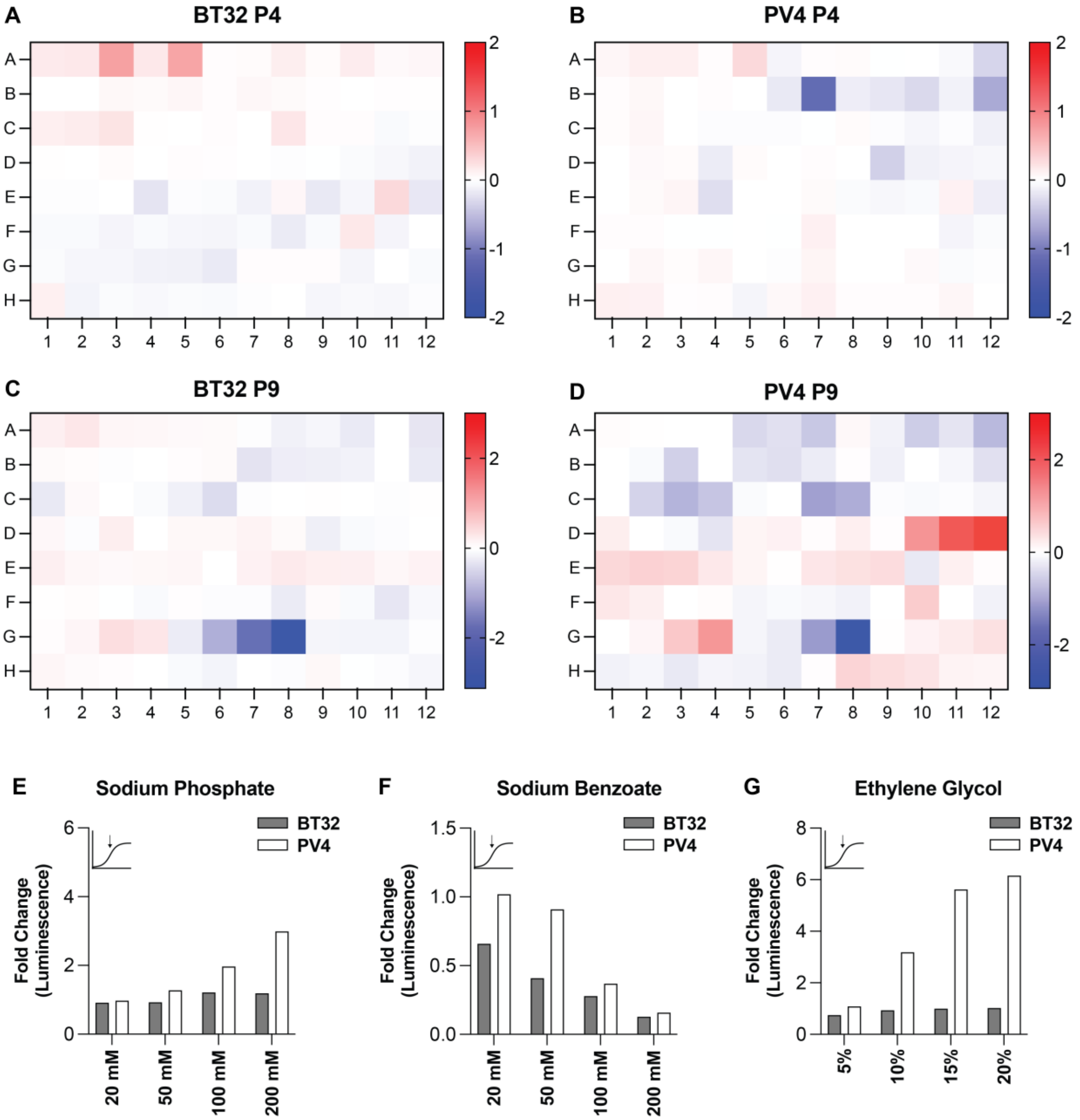


Supplementary Figure 14: Application of Biolog Phenotype Array plates identify few inducing environments for BT32 and PV4 candidate biosensors.

(A) BT32 Fold Change to plate median relative luminescence ($RLU/OD_{600}$) for Biolog Phenotype Array Plate P4 (Phosphorus and Sulfur Utilization Assays).
(B) PV4 Fold Change to plate median relative luminescence ($RLU/OD_{600}$) for Biolog Phenotype Array Plate P4 (Phosphorus and Sulfur Utilization Assays).
(C) BT32 Fold Change to plate median relative luminescence ($RLU/OD_{600}$) for Biolog Phenotype Array Plate P9 (Osmotic Ionic Response Assay)
(D) PV4 Fold Change to plate median relative luminescence ($RLU/OD_{600}$) for Biolog Phenotype Array Plate P9 (Osmotic Ionic Response Assay)

(E) Fold Change of BT32 and PV4 to Sodium Phosphate at increasing concentrations (Plate P9, G1-G4, n=1)
(F) Fold Change of BT32 and PV4 to Sodium Benzoate at increasing concentrations (Plate P9, G5-G8, n=1)
(G) Fold Change of BT32 and PV4 to Ethylene Glycol at increasing concentrations (Plate P9, G9-G12, n=1)

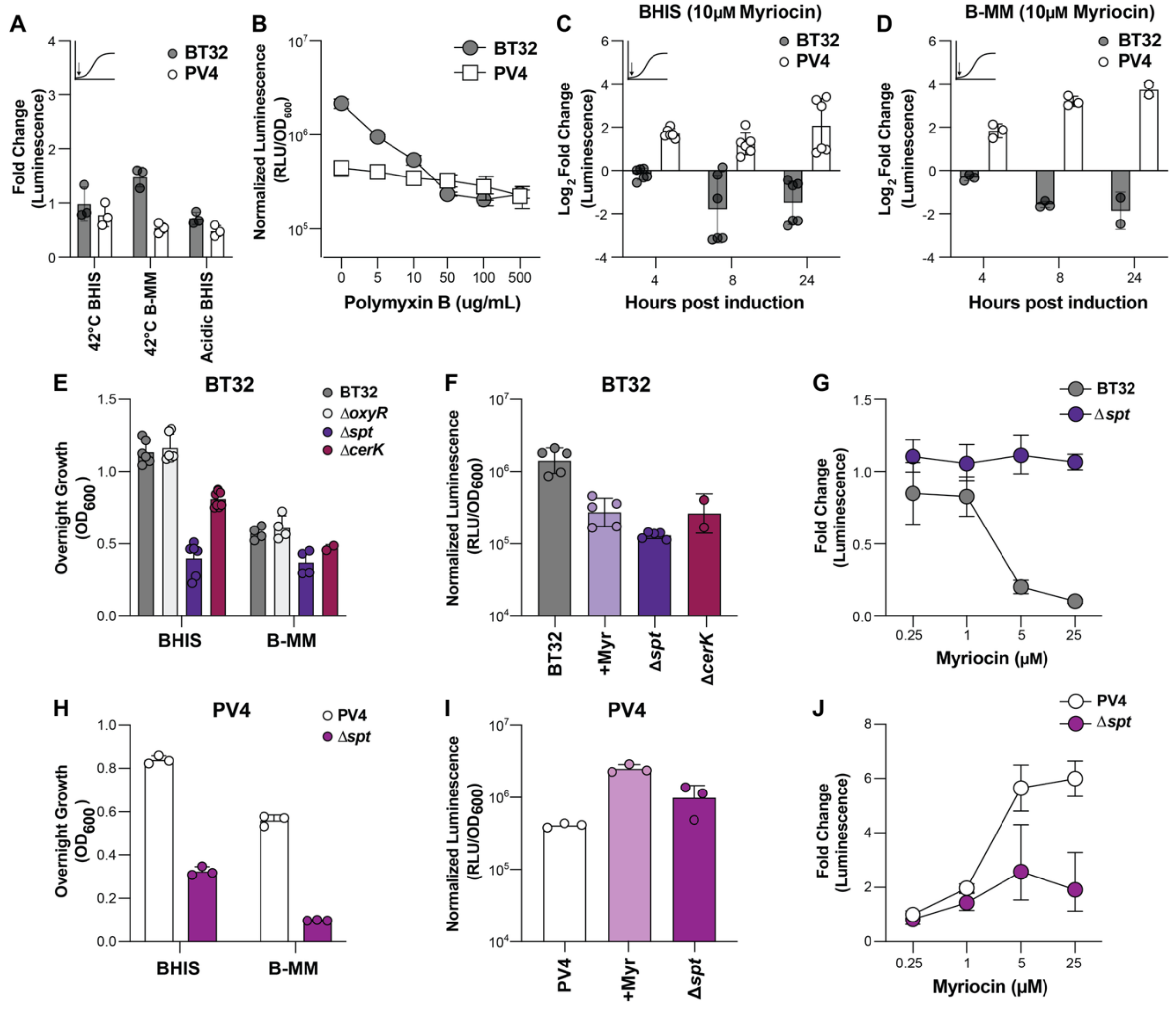


Supplementary Figure 15: BT32 and PV4 do not respond to general stressors heat but do respond to sphingolipid-associated conditions including polymyxin B, myriocin, and genetic deletion of the *spt* (Bt and Pv) and *cerK* (Bt) gene

(A) Subculture in the presence of inducing environment does not induce BT32 or PV4 7-8 hours after subculture and induction. Measured as fold change of relative luminescence of induction vs negative control (equal volume of solvent) (n=3).
(B) Dose-response of BT32 and PV4 to increasing concentrations of Polymyxin B in BHIS media. Cultures were grown to mid-log and induced with equal volumes of Polymyxin B, then measured 2 hours post induction.
(C) Subculture of BT32 and PV4 10 uM Myriocin over time in BHIS media. BT32 and PV4 have divergent responses (Log2FoldChange Relative Luminescence (RLU/OD)) to myriocin. (n=6)
(D) Subculture of BT32 and PV4 10 uM Myriocin over time in BMM media. BT32 and PV4 have divergent responses (Log2FoldChange Relative Luminescence (RLU/OD)) to myriocin. (n=3, 2 for 24 hours)

(E) Overnight growth (OD600) of BT32 and deletion strains Δ*oxyR*, Δ*spt*, and Δ*cerK* in BHIS and BMM media. (n=6 BHIS, 4 BMM)
(F) Normalized Relative Luminescence (RLU/OD600) of BT32, BT32 subcultured with 5 uM myriocin, BT32Δ*spt*, and BT32Δ*cerK* in B-MM 24 hours post subculture and induction.
(G) Dose response to 5 uM myriocin 24 hours post subculture with inducer, measured by fold change to DMSO controls, in BT32 and deletion strain BT32Δ*spt.*
(H) Overnight growth (OD600) of PV4 and deletion strain Δ*spt* in BHIS and BMM media. (n=3 BHIS, 3 BMM)
(I) Normalized Relative Luminescence (RLU/OD600) of PV4, PV4 subcultured with 5 uM myriocin, and PV4Δ*spt*, in B-MM 8 hours post subculture and induction.
(J) Dose response to 5 uM myriocin 8 hours post subculture with inducer, measured by fold change to DMSO controls, in PV4 and deletion strain PV4Δ*spt.*

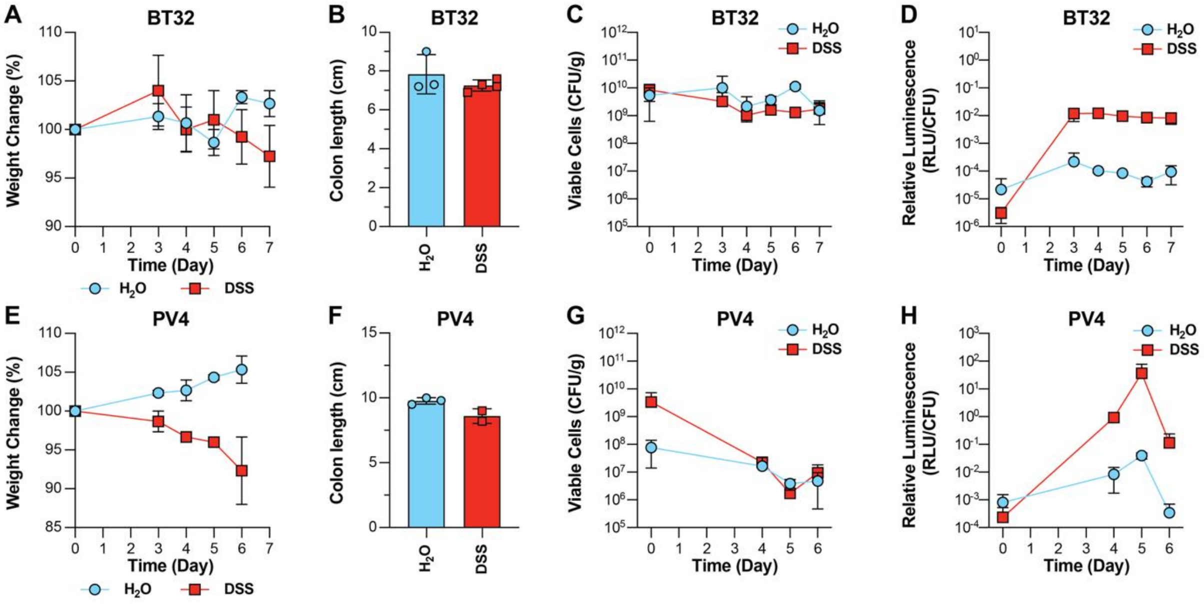


<u>Supplementary Figure 16:</u> BT32 and PV4 biosensors engraft and respond to DSS-induced colitis in gnotobiotic disease model

(A) Percent change in body weight relative to baseline (Day 0) for BT32 engrafted $H_2O$ and DSS conditions (n = 3 $H_2O$, 4 DSS).
(B) Colon length (cm) of BT32 $H_2O$ and DSS groups
(C) Bacterial Load of BT32 (Viable Cells, CFU per g of feces) over experiment time course for $H_2O$ and DSS groups
(D) Relative Luminescence (RLU / CFU) of BT32 biosensor over experiment time course for $H_2O$ and DSS groups
(E) Percent change in body weight relative to baseline (Day 0) for PV4 engrafted $H_2O$ and DSS conditions (n = 3 $H_2O$, 2 DSS).
(F) Colon length (cm) of PV4 $H_2O$ and DSS groups
(G) Bacterial Load of PV4 (Viable Cells, CFU per g of feces) over experiment time course for $H_2O$ and DSS groups
(H) Relative Luminescence (RLU / CFU) of PV4 biosensor over experiment time course for $H_2O$ and DSS groups

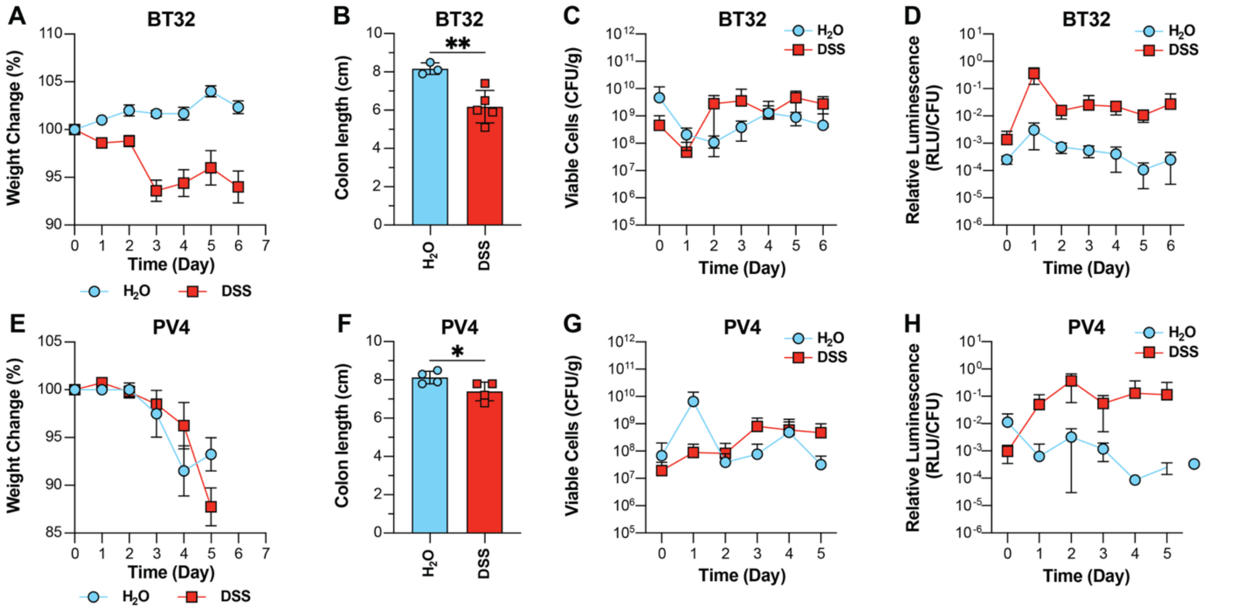


Supplementary Figure 17: BT32 and PV4 biosensors engraft and respond to DSS-induced colitis in conventional disease model

(A) Percent change in body weight relative to baseline (Day 0) for BT32 engrafted $H_2O$ and DSS conditions (n = 3 $H_2O$, 4 DSS).
(B) Colon length (cm) of BT32 $H_2O$ and DSS groups
(C) Bacterial Load of BT32 (Viable Cells, CFU per g of feces) over experiment time course for $H_2O$ and DSS groups
(D) Relative Luminescence (RLU / CFU) of BT32 biosensor over experiment time course for $H_2O$ and DSS groups
(E) Percent change in body weight relative to baseline (Day 0) for PV4 engrafted $H_2O$ and DSS conditions (n = 4 $H_2O$, 4 DSS).
(F) Colon length (cm) of PV4 $H_2O$ and DSS groups
(G) Bacterial Load of PV4 (Viable Cells, CFU per g of feces) over experiment time course for $H_2O$ and DSS groups
(H) Relative Luminescence (RLU / CFU) of PV4 biosensor over experiment time course for $H_2O$ and DSS groups

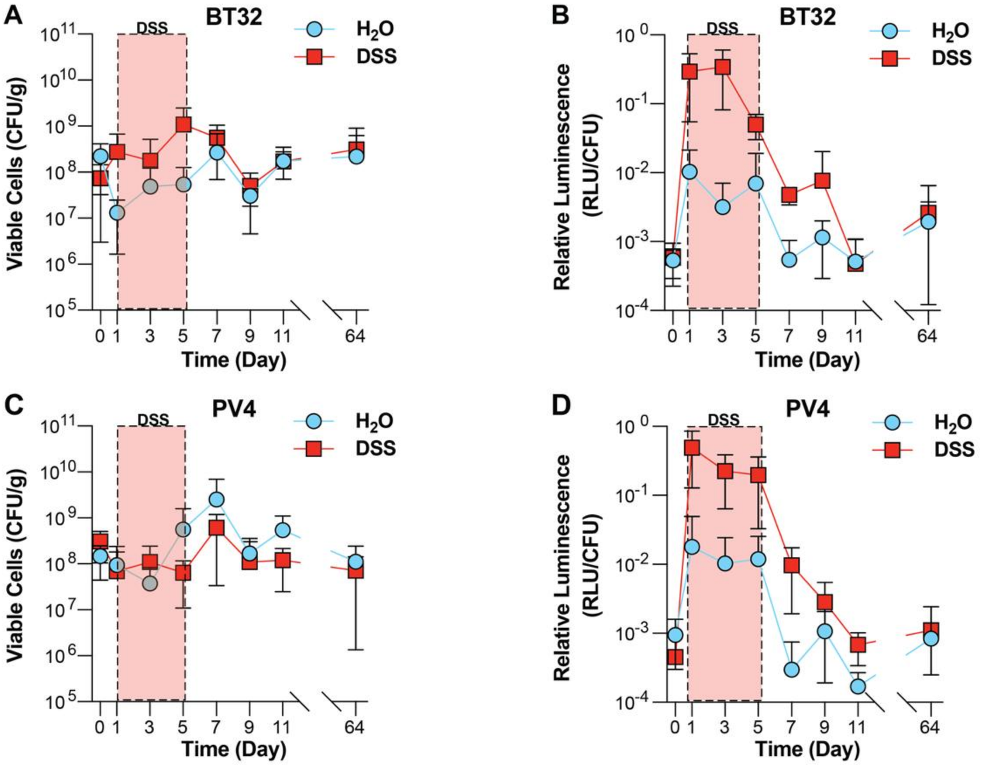


Supplementary Figure 18: BT32 and PV4 biosensors engraft, respond to inflammation and return to baseline in DSS Recovery Model

(A) Bacterial Load of BT32 (Viable Cells, CFU per g of feces) over experiment time course for $H_2O$ and DSS groups. BT32 engrafts for at least 64 days after DSS treatment.
(B) Relative Luminescence (RLU / CFU) of BT32 biosensor over experiment time course for $H_2O$ and DSS groups. Biosensor returns to H2O levels after Host recovery.
(C) Bacterial Load of PV4 (Viable Cells, CFU per g of feces) over experiment time course for $H_2O$ and DSS groups. BT32 engrafts for at least 64 days after DSS treatment.
(D) Relative Luminescence (RLU / CFU) of PV4 biosensor over experiment time course for $H_2O$ and DSS groups. Biosensor returns to H2O levels after Host recovery.
